# Theta rhythmicity governs the timing of behavioral and hippocampal responses in humans specifically during memory-dependent tasks

**DOI:** 10.1101/2020.11.09.374264

**Authors:** Marije ter Wal, Juan Linde Domingo, Julia Lifanov, Frederic Roux, Luca Kolibius, Stephanie Gollwitzer, Johannes Lang, Hajo Hamer, David Rollings, Vijay Sawlani, Ramesh Chelvarajah, Bernhard Staresina, Simon Hanslmayr, Maria Wimber

**Affiliations:** School of Psychology & Centre for Human Brain Health, University of Birmingham, Edgbaston, B15 2TT, Birmingham, UK; Max Planck Institute for Human Development, 14195, Berlin, Germany; Universitätsklinikum Erlangen, 91054, Erlangen, Germany; Queen Elizabeth Hospital Birmingham, Edgbaston, B15 2GW, Birmingham, UK; Institute of Neuroscience & Psychology, University of Glasgow, G12 8QB, Glasgow, UK

**Keywords:** Theta oscillations, memory encoding, memory retrieval, reaction times, hippocampus, intracranial EEG, phase locking

## Abstract

Memory formation and reinstatement are thought to lock to the hippocampal theta rhythm, predicting that encoding and retrieval processes appear rhythmic themselves. Here, we show that rhythmicity can be observed in behavioral responses from memory tasks, where participants indicate, using button presses, the timing of encoding or retrieval of cue-object associative memories. We found no evidence for rhythmicity in button presses for visual tasks using the same stimuli, or for questions about already retrieved objects. The oscillations for correctly remembered trials center in the slow theta frequency range (1-5 Hz), while responses from later forgotten trials do not lock to the behavioral oscillation. Using intracranial EEG recordings, we show that the memory task induces temporally extended phase consistency in hippocampal local field potentials at slow theta frequencies, but only for correctly remembered trials, providing a mechanistic underpinning for the theta oscillations found in behavioral responses.

## 3. Introduction

In everyday life, our brains receive a virtually never-ending stream of information that needs to be stored for future reference or requires integrating with pre-existing knowledge. Extensive research has identified the hippocampus as the hub where encoding and retrieval of information is coordinated (for reviews see: Duncan and Schlichting, 2018; Eichenbaum, 2000; O’Reilly and Norman, 2002; Staresina and Wimber, 2019). Information streams within the hippocampus and between hippocampus and cortex are thought to be orchestrated by the phase of the theta rhythm (Colgin, 2016; Düzel et al., 2010; Hasselmo and Stern, 2014). Here, we ask whether theta oscillations clock responses during memory tasks, producing rhythmicity in behavior.

During memory formation, to-be-encoded information processed by cortical regions is sent to the hippocampus and is subsequently uniquely encoded. Conversely, during retrieval, cues trigger the completion of existing patterns encoded in hippocampus, which elicits full reinstatement of the memory in associated cortical regions. Both memory encoding and retrieval have been associated with changes in prominent oscillatory patterns in hippocampal local field potentials (LFPs). The LFP of rodents is dominated by oscillations in the 4-8 Hz theta frequency band, while a broader low frequency band is apparent in humans, with frequencies in intracranial recordings often peaking between 1-5 Hz during memory tasks (Goyal et al., 2020; Griffiths et al., 2019; Jacobs, 2014; Lega et al., 2012). Several studies have shown that encoding of later-remembered items is accompanied by higher theta power compared to later-forgotten items, both in scalp- (Guderian et al., 2009; Staudigl and Hanslmayr, 2013) and subdural recordings (Kahana et al., 1999; Sederberg et al., 2003), as well as in human hippocampus (Fell et al., 2011; Lega et al., 2012; Lin et al., 2017), but see (Herweg et al., 2020). Similarly, phase-amplitude coupling between theta and gamma oscillations increased during successful encoding (Lega et al., 2016; Mormann et al., 2005; Staudigl and Hanslmayr, 2013). Finally, spiking activity of hippocampal neurons was reported to lock to the LFP at theta frequencies (Jacobs et al., 2007), with stronger locking during successful encoding than for later forgotten items (Rutishauser et al., 2010).

During memory retrieval, theta power increased in cortical areas that are involved in reinstatement (Jacobs et al., 2006) and synchronization between these areas and hippocampus increased at theta frequencies (Anderson et al., 2010; Benchenane et al., 2010; Fujisawa and Buzsáki, 2011; Herweg et al., 2016; Watrous et al., 2013). Intriguingly, recall signals in hippocampus precede reinstatement in cortex by about one theta cycle, suggesting hippocampus and cortex communicate within theta ‘windows’ during memory recall (Staresina and Wimber, 2019).

There is wide-ranging evidence that information in the rodent hippocampus is structured by the phase of the theta rhythm. Sequences of past, current and predicted future locations are phase-coded in theta cycles during movement through a familiar environment (Dragoi and Buzsáki, 2006; Foster and Wilson, 2007; O’Keefe and Recce, 1993; Skaggs et al., 1996), a finding that has been extended to temporal event sequences (Allen et al., 2016; Crivelli-Decker et al., 2018; Heusser et al., 2016; Sanders et al., 2019). In recent human studies, reinstatement of remembered associations was found to be theta-rhythmic (Kerren et al., 2018), and remembered spatial goal locations have been found to be represented at different phases of the theta rhythm (Kunz et al., 2019; Watrous et al., 2018).

Theoretical work by Hasselmo and colleagues proposes that the theta rhythm (Hasselmo et al., 2002) separates new information entering hippocampus from reactivated information. Locking of pattern separation and completion to opposing theta phases avoids destructive interference between new and existing memories. Indeed, strengthening of synaptic connections (long-term potentiation) is more likely to occur around the trough of theta cycles (Hyman et al., 2003; Pavlides et al., 1988), while synaptic depression is more pronounced at the peak (Hyman et al., 2003). In line with these findings, it was shown in rodents that communication of new information from cortex to hippocampus predominantly occurs around the theta trough (Amemiya and Redish, 2018; Colgin et al., 2009; Fernández-Ruiz et al., 2017; Lopes-dos-Santos et al., 2018), while retrieval-related spiking activity in hippocampus is mostly observed around the theta peak (Amemiya and Redish, 2018; Colgin et al., 2009; Fernández-Ruiz et al., 2017; Lopes-dos-Santos et al., 2018). Intracranial recordings from epilepsy patients have suggested similar network dynamics, with entorhinal cortex and hippocampus synchronizing their theta phase during encoding, while hippocampus locked to the downstream subiculum during retrieval (Solomon et al., 2019). Furthermore, optogenetically suppressing neural activity during task-irrelevant phases of the theta oscillation improves performance (Siegle and Wilson, 2014), demonstrating that theta phase has a functional link to memory performance.

Consistent locking of encoding and retrieval processes to the theta rhythm predicts that these processes appear as rhythmic. Rhythmicity might therefore be visible in behavioral markers that depend on long-term memory. To our knowledge, no work has tested for rhythmicity in long-term memory-dependent, overt behavior. However, studies on attentional scanning suggest that oscillatory activity can manifest in behavior (VanRullen, 2016). In both monkey and human, visual attention to a cued location periodically switched towards the periphery, to allow for the prioritization of the cued location while maintaining the flexibility to detect changes in other parts of the visual field. Intriguingly, the effect of attentional scanning was visible in behavioral performance, with the hit rate for detection at the cued location of the participants waxing and waning at a theta rhythm (Busch and VanRullen, 2010; Fiebelkorn et al., 2013, 2018; Helfrich et al., 2018; Landau and Fries, 2012).

Here, we ask whether the theta rhythmicity observed in so many studies in humans and rodents translates into an oscillatory modulation of behavior during memory tasks. We analyze responses from hundreds of participants completing a memory task, in which they pressed buttons to indicate the exact time points at which they formed or recalled an associative memory. We find significant oscillations in both encoding and retrieval button presses, with frequencies matching the previously reported lower theta frequency band (1-5 Hz, Jacobs, 2014; Lega, Jacobs, & Kahana, 2012). No oscillatory signatures were present in button presses from task phases that do not depend on memory. In addition, incorrect trials do not lock to the rhythm identified for correct trials. To establish a mechanism for our behavioral findings, we analyze hippocampal LFPs recorded in epilepsy patients, and find temporally extended phase locking in the low theta range during memory-dependent task phases, which is present for correct but not incorrect trials. Finally, we show that encoding and retrieval trials show maximal phase alignment at opposite phases of the theta rhythm. Together, our results demonstrate that hippocampus-dependent, theta-rhythmic memory processes can be detected in human behavior.

## 4. Results

### 4.1. Button presses indicate the timing of memory-dependent and -independent processing

In this study we asked whether signatures of hippocampal rhythms can be found in behavioral responses during the encoding and retrieval of memories. We analyzed the data from 226 participants who performed one of several related associative memory tasks. The memory tasks consisted of multiple blocks with encoding, distractor and retrieval phases (Figure 1A). During encoding phases, participants were shown a cue (verb or scene image), followed by a stimulus (photo or drawing of an object), and pressed a button when they made an association between cue and stimulus. This provided us with an estimate of the timing of successful memory formation (Encoding button press). During retrieval phases, participants were shown the cues in random order and were asked to remember the associated objects. One group of participants (group 1; n=71) indicated the moment they remembered the object by pressing a button (Retrieval button press) and then answered one or two catch questions (e.g. ‘animate or inanimate?’) about the already reinstated object (Catch-after-retrieval button press). A second group (group 2; n=155) was shown the catch question before the cue appeared. This group thus needed to mentally reinstate the object and press the button as soon as they were able to answer the question (Catch-with-retrieval button press), and hence this button press indicated the time of subjective memory retrieval in this group. Each participant memorized between 64 and 128 cue-object pairs. Objects, cues and catch questions varied between experiments; for details see Methods and Table S1.

**Figure 1.**
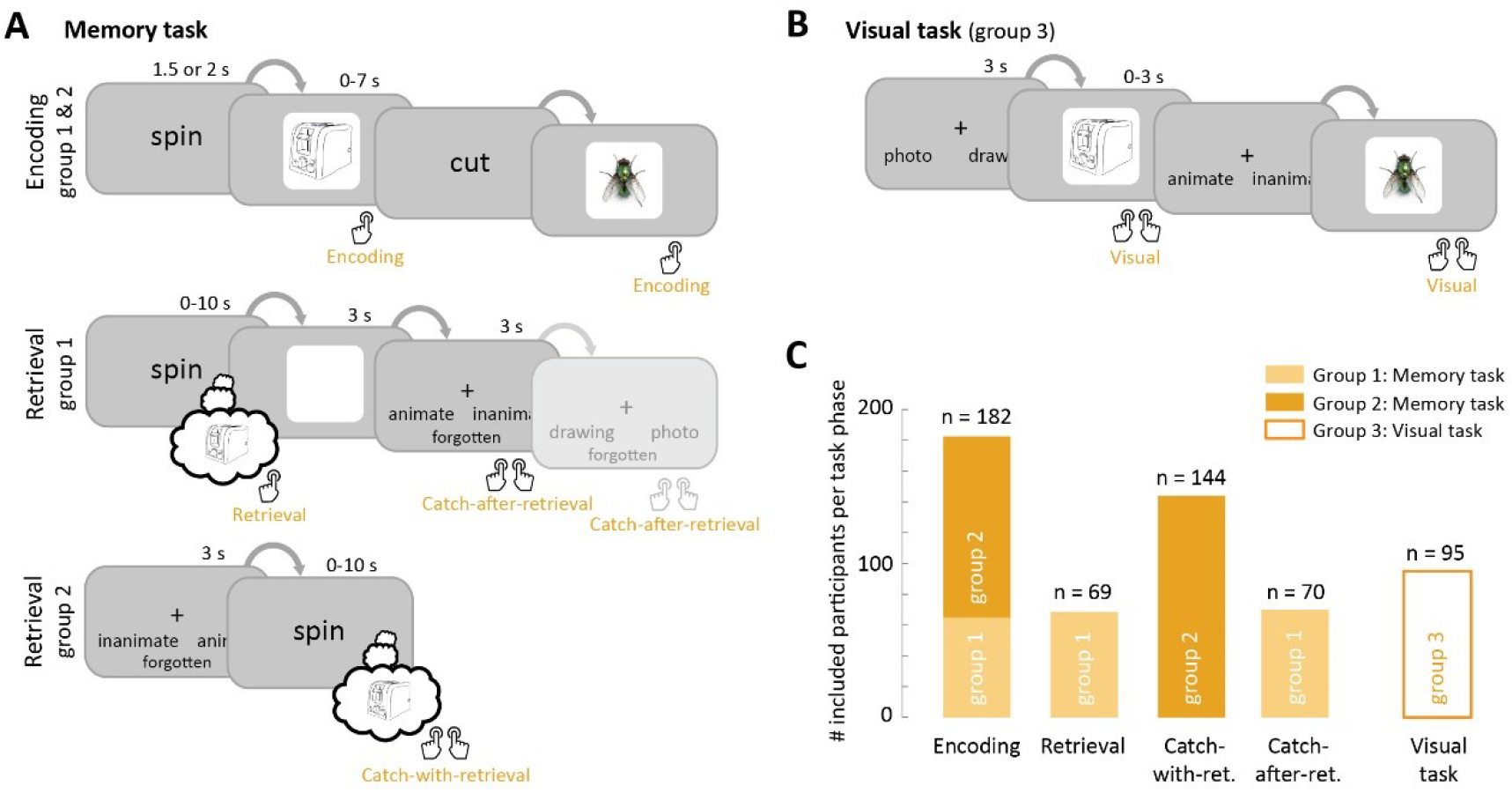
Button presses indicate the timing of memory-dependent and -independent processing. **A**: Structure of the memory task. Two groups of participants (group 1 & 2; n=226) completed blocks consisting of an encoding phase (top row) in which they associated cues (‘spin’, ‘cut’) to objects, a distractor phase (not shown), and one of two versions of a retrieval phase (bottom rows), in which they answered catch questions about remembered objects (‘animate or inanimate?’, ‘photo or drawing?’); **B**: Structure of the visual task. A separate group of participants (group 3; n=95) answered questions about objects on the screen, using the same questions and stimuli as the memory task; **C**: Number of participants that were included in further analyses, after exclusion of participants with a high number of incorrect and/or timed-out trials (Figure S1A). Note that participants in group 1 contributed button presses to 3 task phases (Encoding, Retrieval and Catch-after-retrieval), group 2 contributed to 2 (Encoding and Catch-with-retrieval) and group 3 to 1 task phase (Visual).

In order to separate memory processes from perceptual elements of the task, we asked a separate group (group 3; n=95) to perform visual control tasks (Figure 1B). Participants were shown a question followed by an object, and answered the question by pressing a button (Visual button press). The same questions and objects were used as for the memory tasks. Note that the button presses from the visual task do not depend on episodic memory, as they pertain to objects that are constantly visible. Answers to the catch question for memory group 1 (Catch-after-retrieval button press) are also not expected to rely on hippocampal memory retrieval, since they are asked after objects are already (more or less fully) reinstated. Note that answering the catch questions after retrieval is, however, likely to rely on maintenance of the successfully retrieved object in working memory. In this manuscript we use the term memory-dependent as relying on hippocampal dependent long-term memory.

We analyzed the performance of the participants based on the catch questions. Participants who performed at chance level, as assessed by a binomial test, were excluded from further analyses (n=12 for memory task; n=0 for visual task, see Figure S1A). In addition, 32 (1) participants with sufficient memory performance had a low trial count for the encoding (retrieval) phase due to trial time-outs, and as a result were excluded for encoding (retrieval) phase analyses (Figure S1A). The remaining participants (Figure 1C) performed the tasks well, with an average performance of 84.0% (range 56.3-100%) for the memory groups and 96.3% (range 78.2-100%) for the visual group (Figure S1B). For the number of responses and reaction times per task phase see Figure S1 and Table S3.

### 4.2. Oscillatory patterns can be detected in behavioral responses using the O-score

The button presses from the memory tasks provided us with estimates of when participants formed and reinstated memories on each trial. In Figure 2A the button presses from all retrieval trials of one example participant are shown, as well as the smoothed response density across trials. We asked whether the response densities showed oscillations, as suggested by the trend-removed trace in Figure 2A (right), and whether these patterns differed between memory-dependent and -independent task phases. To address this we used the Oscillation score (Muresan et al., 2008), a method developed to assess oscillations in spike trains. This procedure identifies the dominant frequency in the response time stamps, and provides a normalized amplitude at this frequency: the O-score. We computed O-scores for correct responses per participant and task phase.

**Figure 2.**
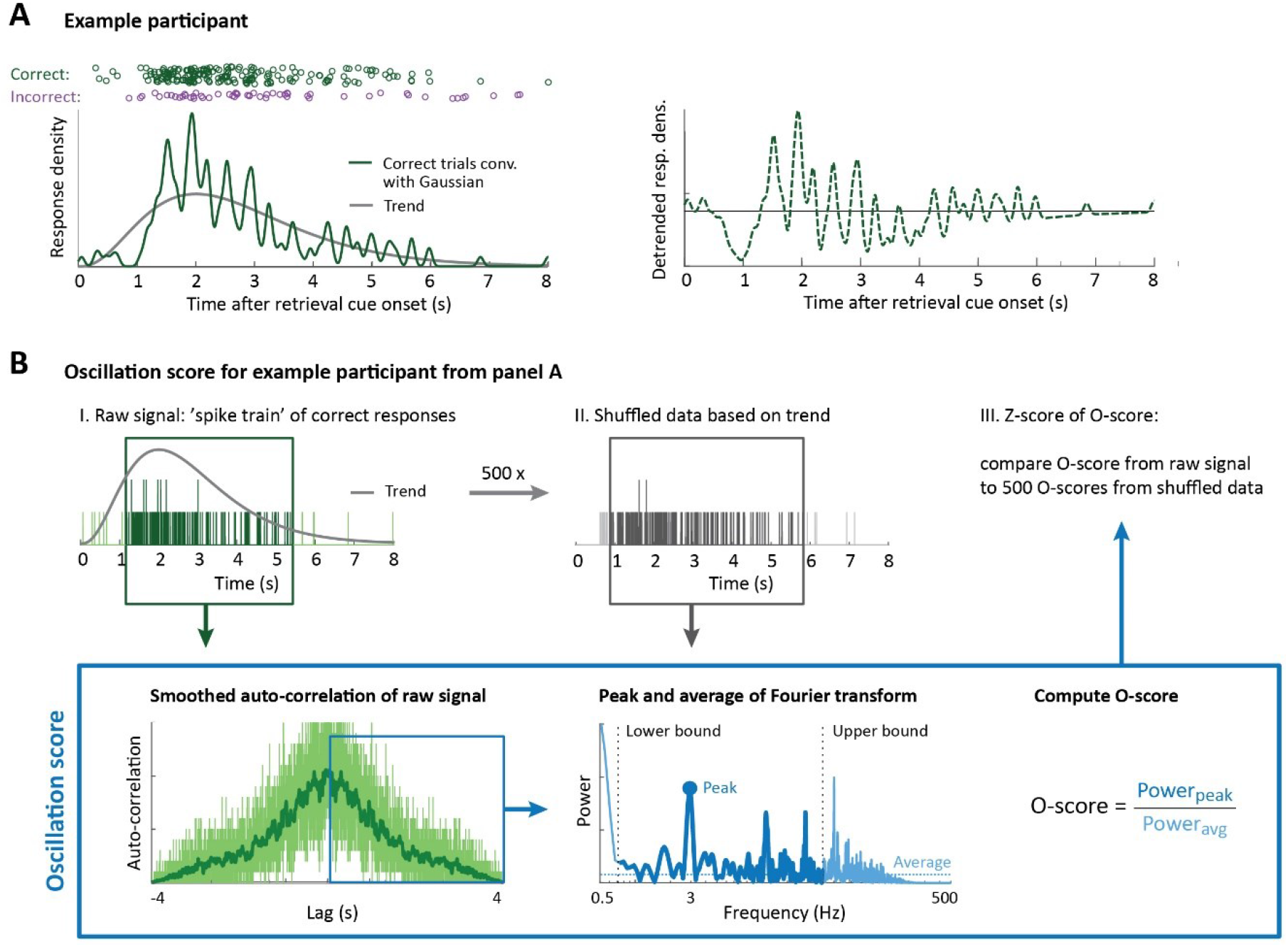
Oscillatory patterns can be detected in behavioral responses using the O-score. **A**: Timing of retrieval button presses for an example participant. Each circle is one button press, with correct trials in green and incorrect trials in purple. Convolving the correct trials with a Gaussian kernel (left panel, solid green line) reveals an overall trend in response density (left panel, grey line) as well as an oscillatory modulation (right panel, dashed green line); **B**: Step-by-step representation of the Oscillation score method, for the participant from panel A. O-scores were computed for the original data (dark green) and 500 reference datasets with the same overall response trend (grey) following the procedure from (Muresan et al., 2008), summarized in the blue box. For details see main text and Methods.

In brief, after removal of early and late outliers (Figure 2B, step I), we computed the O-score as follows (Figure 2B, blue box): The auto-correlation histogram (ACH) was computed for the button presses from correct trials and smoothed with a Gaussian kernel (σ=2 ms) to reduce noise. Then the central peak of the ACH was removed. All remaining positive lags were Fourier transformed and, within a frequency range of interest (adjusted per participant based on the signal length (lower bound) and number of responses (upper bound), with a minimum of 0.5 Hz and maximum of 40 Hz), the frequency with the highest magnitude was found. The O-score was computed by dividing the peak magnitude by the average of the entire spectrum.

The O-score indicates how much the peak frequency stands out against other frequencies, but does not take into account the overall response structure in the button presses (grey trend curves in Figure 2) and the limited number of data points, which could introduce a frequency bias. To account for this, we fitted a trend curve for each participant and generated 500 random series of button presses based on this structure, with the same number of data points as the original dataset (Figure 2B, II). We then computed the O-score for the random datasets at the peak frequency from the intact data and Z-transformed the original O-score against the 500 reference O-scores (Figure 2B, III). This allowed us to statistically assess the oscillatory strength for each participant and task phase, as well as perform second level statistical assessments across participants. We validated the performance of the O-score and Z-scoring methods using simulated data (Supplementary Results and Figure S5).

### 4.3. Behavioral responses oscillate at theta frequencies for memory-dependent task phases

O-scores were high for encoding and retrieval button presses from both versions of the memory task (Figure 3A). Each of the memory-dependent task phases reached significance across participants, specifically Encoding (t(181)=6.20; p<0.0001), Retrieval (t(68)=4.58; p<0.0001) and Catch-with-retrieval, (t(143)=5.08; p<0.0001; all Bonferroni-corrected for 5 comparisons; effect sizes in Table S4). In addition, the proportion of participants with significant O-scores was high (Figure 3B): 74.7% for Encoding, 69.6% for Retrieval and 76.4% for Catch-with-retrieval. On the other hand, no evidence for a behavioral oscillation was found for memory-independent task phases, as O-scores for the catch questions after memory reinstatement and the visual task did not reach significance across participants (Catch-after-retrieval: t(69)=1.69; p=0.24; Visual: t(94)=−4.10; p=1; Bonferroni-corrected for 5 comparisons; Note that the t-value captures deviation from the reference-defined threshold, hence both non-significant and negative t-values signify lack of evidence for oscillations). For these task phases the proportion of participants with significant O-scores was lower than for memory-dependent phases: 64.3% for Catch-after-retrieval and 37.9% for the Visual task. Raw O-scores (i.e. without Z-scoring) showed a similar pattern across task phases (Figure S3A).

**Figure 3.**
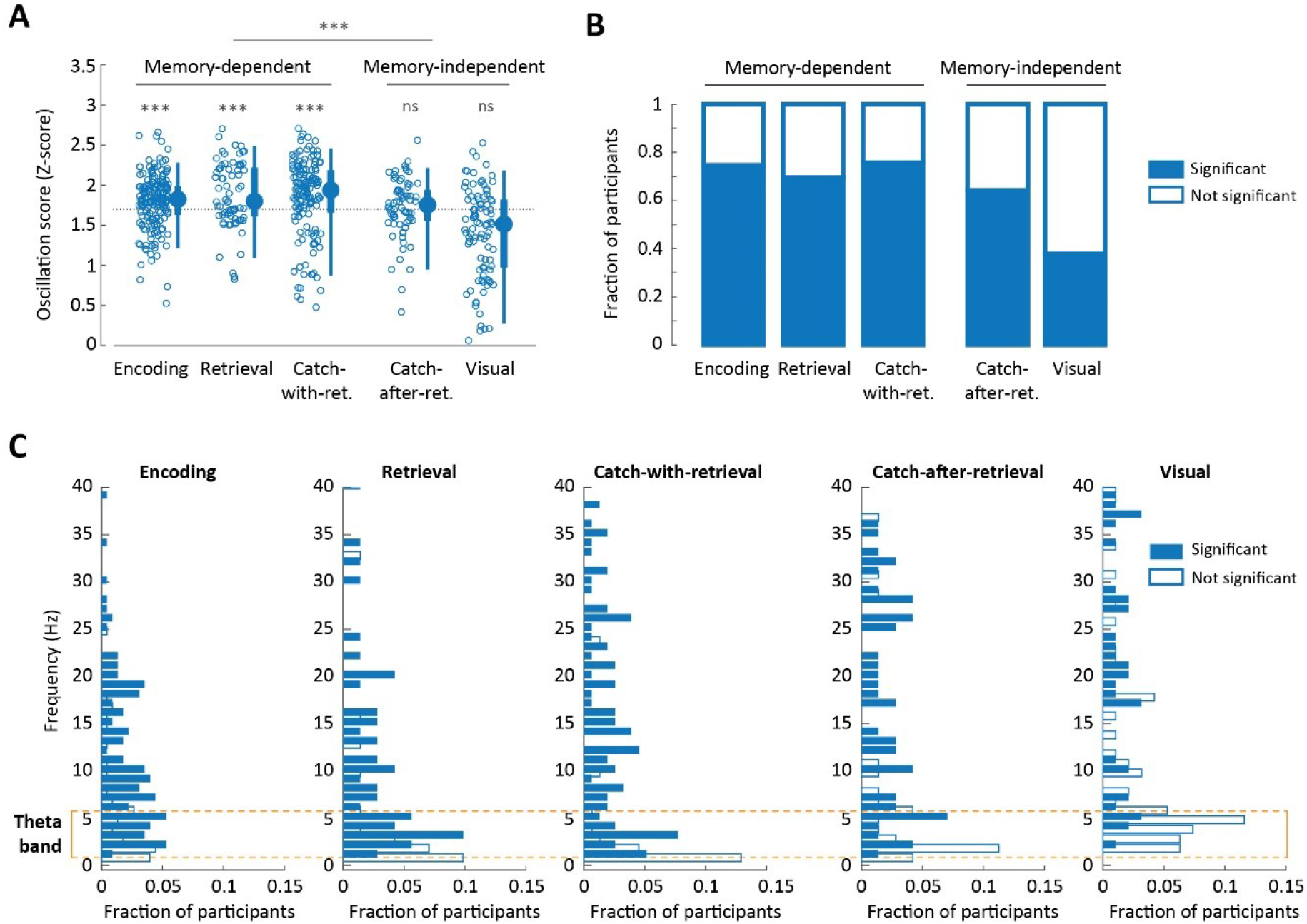
Behavioral responses oscillate at theta frequencies for memory-dependent task phases. **A**: Scatter plot of O-scores (Z-scored) per task phase, where each circle is one participant, and box plots representing the 5, 25, 50, 75 and 95% bounds of the O-score distribution across participants. The dashed line gives the significance threshold for single participants (α=0.05, one-tailed, Z-distribution). The outcome of a second level t-test is given above each task phase, and the comparison between memory-dependent and -independent task phases is given at the top (linear mixed model, see main text and Methods). See Figure S3 for raw O-scores and additional statistics; ns: not significant; *: 0.05≥p>0.01; **: 0.001≥p>0.001; ***: p≤0.0001, Bonferroni-corrected for 5 comparisons; **B**: Proportion of participants with significant (i.e. above the Z-threshold in A; blue bars) and non-significant O-score (white bars); **C**: Histograms of peak frequencies per task phase, for participants with significant (blue bars) and non-significant O-scores (white bars). The yellow outline indicates the 1-5 Hz frequency band. Participant numbers can be found in Figure 1C.

To test whether memory-dependent task phases had significantly higher O-scores than memory-independent phases across participants, we fitted a linear mixed-effects model to the Z-scored O-scores. Fixed terms in this model were memory dependence and the length of the time series, which varied substantially between task phases (Figure S1C); we included the intercept per subject as random effect, to address potential dependencies due to the fact that participants of the memory task contributed 2 or 3 data points. We found strong support for an effect of memory dependency on O-score, with significantly higher Z-scored O-scores for memory-dependent than memory-independent task phases (Figure 3A; coefficient=0.28; 95% CI: 0.20-0.36; t(556)=6.55; p<0.0001). This was unaffected by time series length (coefficient=0.0022; 95% CI: −0.0035-0.0080; t(556)=0.77; p=0.44). Additional paired t-tests (Figure S3 B and C, effect sizes in Table S4) showed that O-scores for Catch-after-retrieval were lower than for Retrieval from the same participants (two-tailed paired t-test; t(68)=2.46; p=0.0495; Encoding versus Catch-after-retrieval did not reach significance: t(64)=1.98; p=0.16; Bonferroni-corrected for 3 comparisons), while Encoding did not differ from Retrieval (t(63)=−0.76; p=1; Bonferroni-corrected for 3 comparisons) and Catch-with-retrieval (t(116)=−0.94; p=1). Furthermore, O-scores were significantly lower for the Visual task than all other task phases (Figure S3; two-tailed two-sample t-tests; Encoding versus Visual: t(274)=7.23; p<0.0001; Retrieval versus Visual: t(161)=5.83; p<0.0001; Catch-with-retrieval versus Visual: t(236)=6.48; p<0.0001; Catch-after-retrieval versus Visual: t(162)=4.04; p=0.0034; Bonferroni-corrected for 4 comparisons).

High O-scores give a strong indication of rhythmic behavior, but for this rhythmicity to be physiologically relevant, we expect some consistency in peak frequency across participants. Indeed, for memory-dependent task phases most participants showed peak frequencies (Figure 3C) between 1 and 5 Hz or harmonics of this range, with 31.6%, 41.7% and 27.5% of significant O-scores falling in the 1-5 Hz range for Encoding, Retrieval and Catch-with-retrieval task phases, respectively. These frequencies align with the low theta band identified in recordings from human hippocampus during memory tasks (Jacobs, 2014; Lega et al., 2012). On the other hand, peak frequencies were broadly distributed for Catch-after-retrieval and the Visual task. A linear mixed model fitting the frequencies of significant O-scores, with memory dependency and time series length as fixed effects, and participant ID as random effect, revealed that frequencies for memory-dependent task phases were significantly lower than for memory-independent tasks (coefficient= −5.09; 95% CI: −7.52-−2.67; t(371)=6.55; p<0.0001, post-hoc tests in Table S5). There was a small effect of time series length on frequency (coefficient=−0.20; 95% CI: −0.36--0.046; t(371)=−2.55; p=0.011), with the higher frequencies for memory-independence corresponding to shorter time series. To ensure that the difference in O-scores and frequency between memory dependent and -independent task phases was not caused by the lower frequency limit in the O-score procedure, which required the oscillation period to be no wider than 1/3 of the time series length, we loosened this bound to 2 times the time series length and recomputed the O-scores. This produced similar results (Figure S3E; Catch-after-retrieval: t(69)=0.91; p=0.90; Visual: t(94)=−6.20; p=1; Bonferroni-corrected for 5 comparisons), reaffirming that the identified difference between memory-dependent and -independent task phases is not caused by differences in response times.

### 4.4. Reaction times of incorrect trials are not locked to the behavioral oscillation

The O-scores reported in Figure 3 were based on correct trials only. Due to a low number of incorrect trials, it was not possible to establish whether incorrect trials show oscillatory modulation. However, we were able to test whether incorrect trials locked to the oscillation of the correct trials (correcting for fitting bias, see below) for every participant with a significant O-score. The instantaneous phase of the oscillation was determined by smoothing and filtering the correct response trace around the participant’s peak frequency (for an example see Figure 2A, solid green line) and performing a Hilbert transform. We then determined the phases at which the incorrect button presses occurred (Figure 4, purple lines). Similarly, we found the phase of each correct response relative to all other correct trials, by recomputing the instantaneous phase without the trial of interest (avoiding circularity), repeating this for every correct trial (Figure 4, green). As expected, for all memory-dependent task phases, correct trials were more likely to occur around the peak of the oscillation, assessed by a V-test for non-uniformity around 0 degrees (Encoding: V(11398)=1861.1; p<0.0001; Retrieval: V(8036)=1174.3; p<0.0001; Catch-with-retrieval: V(23815)=2847.5; p<0.0001; Bonferroni-corrected for 3 comparisons; Note that the high trial count can inflate test results). On the other hand, phase distributions were uniform for incorrect trials (Encoding: V(1270)=44.1; p=0.12; Retrieval: V(954)=11.2; p=0.91; Catch-with-retrieval: V(5328)=73.8; p=0.23; Bonferroni-corrected for 3 comparisons). This suggest that incorrect trials did not lock to the rhythm of the correct trials, while correct responses did lock to the oscillation from other correct trials. We did not perform this analysis for memory-independent task phases, as we found no evidence for oscillations in correct responses for these phases.

**Figure 4.**
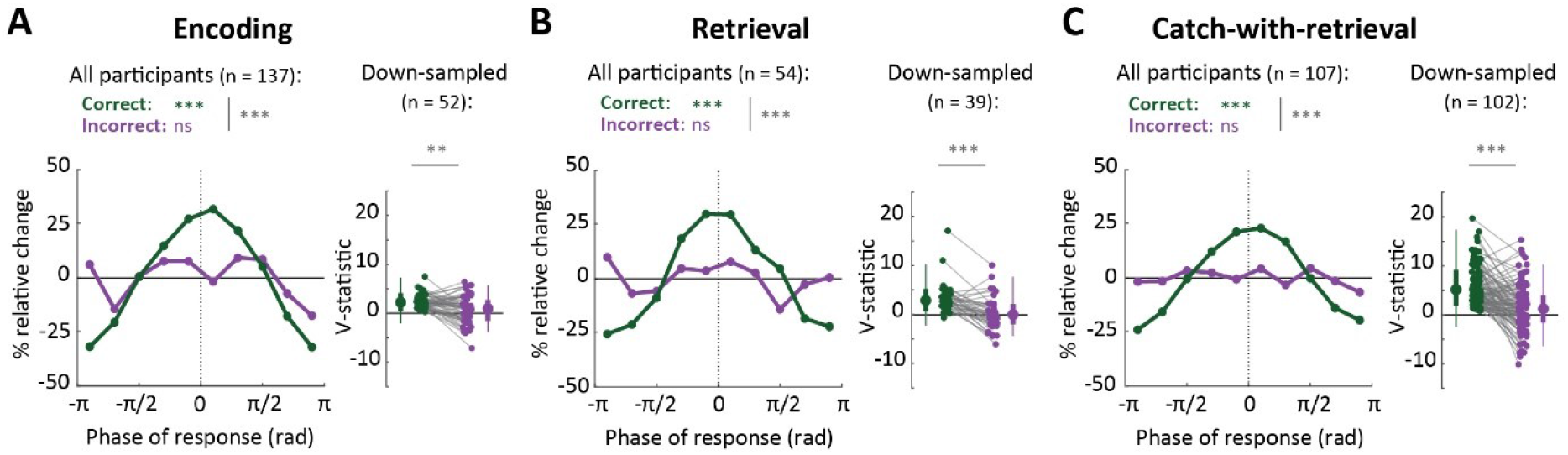
Reaction times of incorrect trials are not locked to the behavioral oscillation. Phase distributions of incorrect responses relative to all correct responses (purple) and of correct responses relative to all other correct responses (green) for the task phases with significant O-scores: Encoding (**A**), Retrieval (**B**) and Catch-with-retrieval (**C**). The left panels shown deviations from uniform phase distributions across all participants with significant O-scores. Statistics for correct and incorrect trials individually were obtained with a V-test for non-uniformity of the distribution around phase 0, and a permutation test was used to compare correct with incorrect distributions to a trial label-shuffled reference distribution (500 permutations, see Methods). Right panels show V-statistics for correct and incorrect trials of participants with at least 10 incorrect trials (each grey line is one participant), after down-sampling the number of correct trials to the number of incorrect trials. Shown are the mean V-statistics per participant (dots) and the distribution of V-statistics across all participants (box plots indicating the 5, 25, 50, 75 and 95% bounds of the distributions), which were compared with a two-tailed paired t-test. ns: not significant; *: 0.01<p<0.05; **: 0.0001<p<0.01; ***: p<0.0001, Bonferroni-corrected for 3 comparisons.

We next directly tested whether there was a difference in phase modulation between correct and incorrect trials, while accounting for potential biases caused by differences in trial count and procedure. We shuffled the correct and incorrect trial labels 500 times for every participant (i.e. keeping the original trial counts and response times), and computed the V-statistics of shuffled-correct and shuffled-incorrect trials as described previously. Stronger phase modulation for correct than for incorrect trials should result in a bigger difference in V-statistics for the real-correct and real-incorrect trials than for the corresponding label-shuffled trials. Indeed, phase modulation was significantly stronger for correct than for incorrect trials across participants (Encoding: p<0.002; Retrieval: p<0.002; Catch-with-retrieval: p<0.002; 500 permutations).

For participants with a sufficient number of incorrect trials (at least 10) we also compensated for any biases by subsampling the number of correct trials to the number of incorrect trials (repeated 100 times), and recomputing the phases of both the selected correct and the incorrect trials relative to the remaining correct trials (Figure 4, right panels). For these participants, the subsampling procedure also demonstrated significantly higher phase modulation for correct trials than for incorrect trials for each of the memory-dependent task phases (two-tailed paired t-test; Encoding: t(51)=4.07; p=0.00049; 95% CI: <0.0001–0.14; Retrieval: t(38)=5.41; p<0.0001; 95% CI: <0.0001-0.024; Catch-with-retrieval: t(101)=7.25; p<0.0001; 95% CI: <0.0001; Bonferroni-corrected for 3 comparisons). In conclusion, all comparisons demonstrate that correct responses show substantially more phase locking than incorrect trials. Combining these findings with our previous analyses, our data suggest that correct trials show substantial behavioral oscillations, but that incorrect trials do not lock to this oscillation.

### 4.5 Increased phase locking of hippocampal local field potentials during encoding and retrieval

The data reported so far indicate that across different trials, memory-relevant behavioral responses fall onto a consistent phase of a theta oscillation. The presence of such an oscillation, determined on the basis of one response per trial, implies phase consistency across trials in the neural oscillations underlying the formation and reinstatement of memory. We hypothesized that this phase consistency is induced by events in the trial, namely by the onset of the stimulus or cue, and persists until the participant successfully encodes or retrieves the memory (expected to slightly precede the button press). Given the theta peak frequency identified in the behavioral data and the nature of the task, we expected this phase consistency to occur in the hippocampus. To test these predictions, we recorded hippocampal LFPs in 10 epilepsy patients who were undergoing seizure monitoring using intracranial EEG. These patients performed the same memory task as healthy participants and their behavioral data are included in the results in Figures 1, 3 and 4. We recorded from 42 Behnke-Fried micro electrodes located in hippocampus, as well as 3 electrodes in parahippocampal cortex and amygdala (Figure 5A). Only data from the hippocampal electrodes are included here.

**Figure 5.**
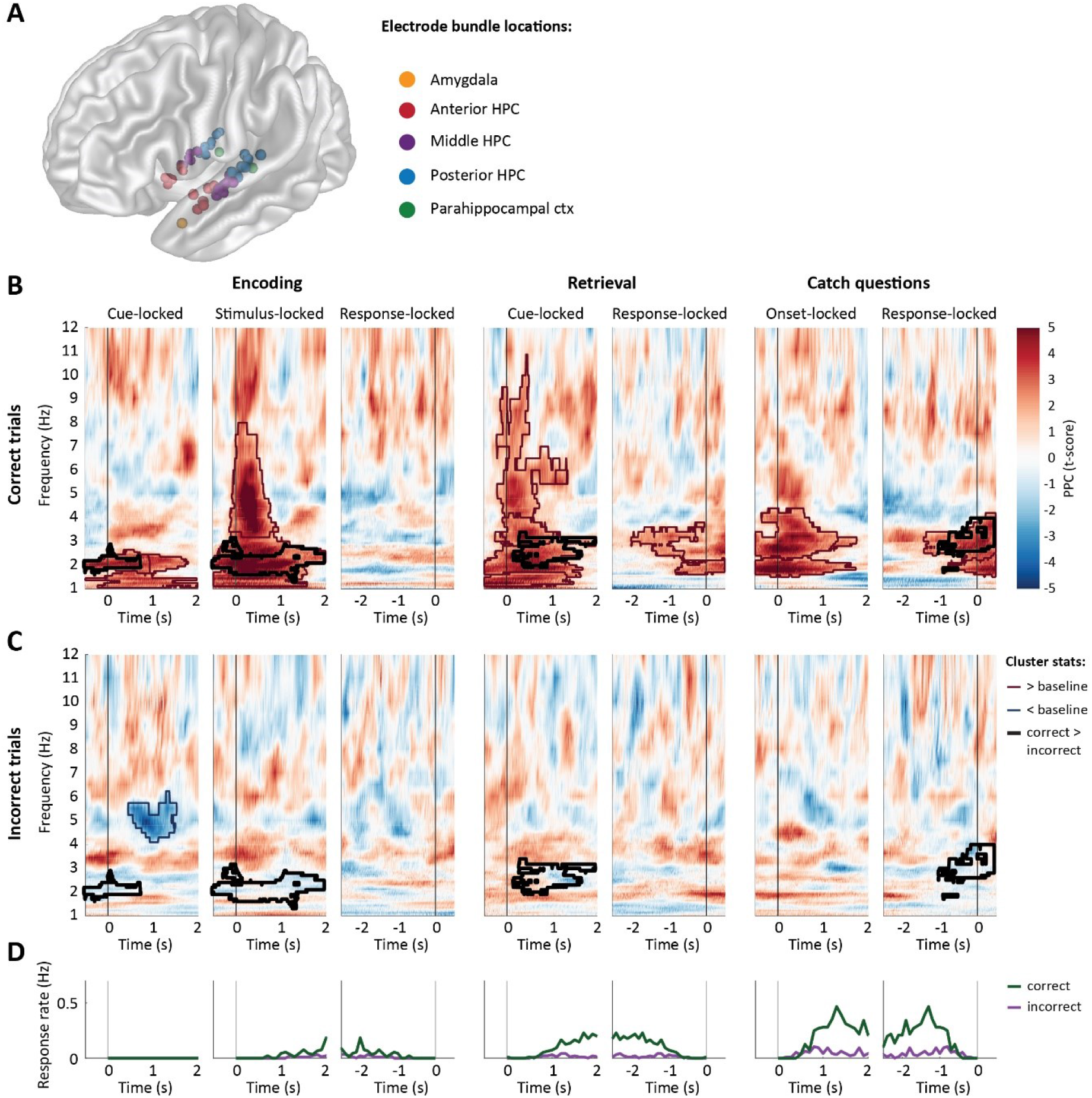
Increased phase locking of hippocampal local field potentials during encoding and retrieval. **A**: Locations of iEEG electrode bundles (n=45), across all 10 participants, color-coded to indicate 5 regions of interest (yellow: amygdala; red: anterior hippocampus (HPC); purple: middle HPC; blue: posterior HPC; green: parahippocampal cortex); **B & C**: Pairwise Phase Consistency (PPC, color-coded, second level t-score) between correct (B) and incorrect (C) trials, locked to cue/stimulus onset or response of encoding (left column), retrieval (middle) and catch trials (right). Significant changes from baseline (α=0.05, permutation test against time-shuffled trials) are indicated separately for increases (red) and decreases (blue). Black outlines indicate significant differences between correct and incorrect trials (α=0.05, permutation test against shuffled trial labels, see also Figure S4A). For raw PPC values see Figure S4B and for PPCs per patient see Figure S4E&F. **D**: Response rate for correct (green) and incorrect (purple) trials, in the time windows and task phases corresponding to B & C.

To study the phase structure in our hippocampal recordings, we wavelet transformed the LFPs and computed the Pairwise Phase Consistency (PPC; Vinck, van Wingerden, Womelsdorf, Fries, & Pennartz, 2010) across trials for every frequency and time point. The PPC quantifies how similar the LFP phases are across trials. We performed this analysis separately for cue-, stimulus- and response-locked data and normalized the data against the pre-cue baseline for every electrode, before pooling across electrodes and participants. The resulting second level t-scores are shown for correct trials in Figure 5B and for incorrect trials in Figure 5C.

In line with our predictions, we observed a significant (α=0.05) increase in PPC across trials for correct trials after stimulus onset for encoding and after cue onset for retrieval trials, assessed through a cluster-based permutation test against 100 time-shuffled datasets (Maris and Oostenveld, 2007). The clusters of significantly increased PPC (red outlines in Figure 5B) covered a range of frequencies shortly after cue/stimulus onset, but extended in time in a frequency band between 2 and 3 Hz, generally aligning with the frequencies found in the behavioral data of these patients (see Figure S4 J&K). This lower theta cluster lasted up to the response (Figure 5D), and resulted in a significant response-locked PPC cluster for retrieval (response-locked data for encoding showed increased PPC, but did not reach significance). PPC increases were also visible in the raw data (Figure S4B) and appeared as theta oscillations in event-related potentials of correct trials (Figure S4D), confirming that these effects were not caused by changes in the baseline. The increases in PPC were accompanied by increased power for retrieval trials, but not for encoding and catch questions (see Figure S4C), in line with (Vivekananda et al., 2019), suggesting that power and phase were modulated independently.

Interestingly, in line with the behavioral data, we found no significant increases in PPC for incorrect trials, neither for encoding nor retrieval. When we compared the PPC for correct and incorrect trials within electrodes using a cluster-based permutation test against 100 trial-shuffled reference data sets (keeping the number of shuffled-correct and shuffled-incorrect trials intact), we found that the increase in PPC after cue/stimulus onset in the 2-3 Hz frequency band was significantly stronger for correct than for incorrect trials (α=0.05; black outlines in Figures 5B and C). The intracranial data therefore support our hypothesis that memory-relevant task events induce phase resets in the hippocampal theta rhythm that result in persistent phase consistency across trials. The absence of phase consistency in LFPs from incorrect trials supports our behavioral findings that incorrect trials do not lock to the theta rhythm.

### 4.6. Encoding and retrieval occur at different phases of the theta rhythm

The identification of phase consistency across encoding and retrieval trials allows us to determine whether their dominant phases differ, as has been suggested by (Hasselmo et al., 2002). To this end, we identified the time point and frequency at which PPC was maximal for both stimulus- and response-locked trials during encoding, and cue-locked and response-locked trials for retrieval, for each patient. We then computed the phase differences between encoding and retrieval at the corresponding frequencies and time points for every electrode. Indeed, phase differences between stimulus-locked encoding and cue-locked retrieval trials were non-uniformly distributed around 250.7 ± 14.1 degrees (Rayleigh’s Z = 30.87; p < 0.0001). Similarly, phases differed consistently between response-locked encoding and retrieval trials with an average phase difference of 162.4 ± 30.6 degrees (Rayleigh’s Z = 7.35; p = 0.001). Both analyses provided support for a half-cycle difference between encoding and retrieval (V-test around 180 degrees; cue/stimulus-locked: V=33.1; p=0.0047; response-locked: V=46.7; p=0.0013; n=326). We tested whether the observed non-uniformity around 180 degrees phase difference was expected by comparing the V-statistics to those from 500 time-shuffled datasets. We conclude that phase opposition for the response-locked trials was unlikely to be obtained by chance (p=0.042), while for stimulus/cue-locked data (p=0.13), the observed V-statistic could, in part, be inflated by a phase bias, for example due to asymmetry in the theta cycles (Cole and Voytek, 2017).

To test whether phase opposition generalized beyond the time and frequency with the highest PPC, we computed response-locked event-related potentials for each hippocampal electrode bundle, and filtered these in the theta band (Figure 6C). We compared the phase of encoding and retrieval ERPs using V-tests in 200 ms sliding windows. After FDR-correction, 55.8% of tested windows supported phase opposition between encoding and retrieval, a pattern that is unlikely to be produced by chance. Together, these results support both theoretical and empirical findings from previous studies that encoding and retrieval processes occur at different phases of the hippocampal theta rhythm, and generalize these findings to LFP recordings from the human hippocampus.

**Figure 6.**
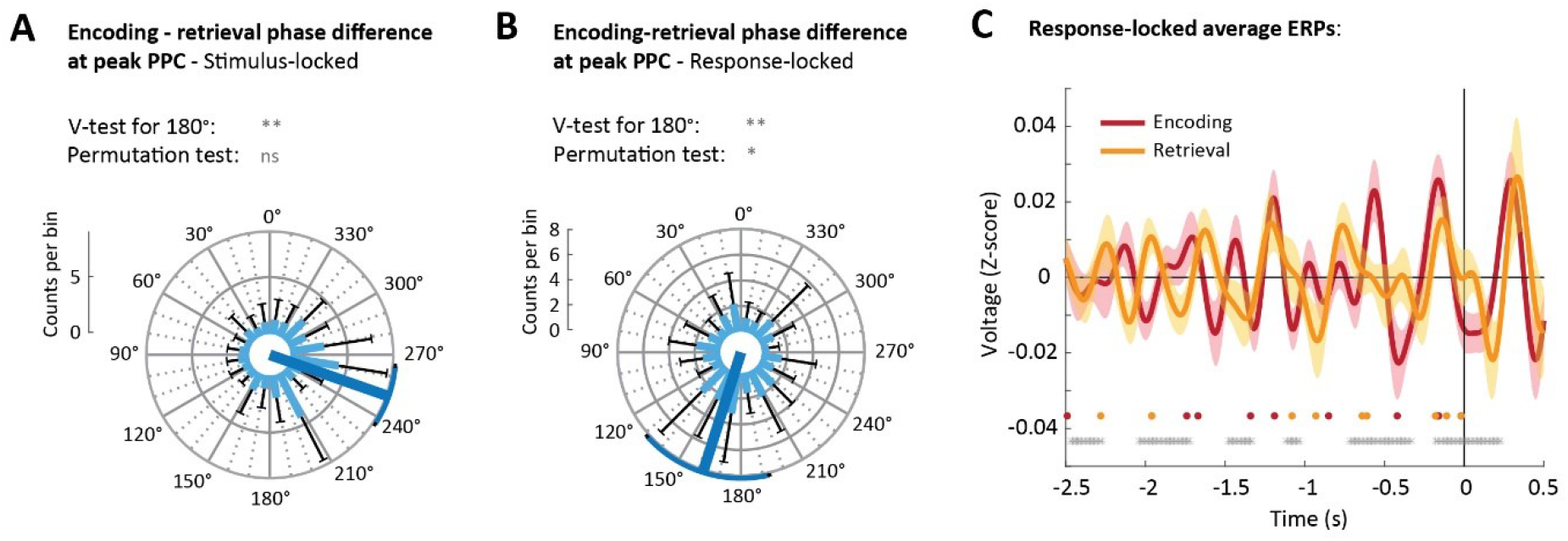
Encoding and retrieval occur at different phases of the theta rhythm. **A & B:** Circular histogram of phase differences between encoding and retrieval at the time and frequency of maximal PPC (A) following stimulus (encoding) and cue onset (retrieval), and (B) before response, across all hippocampal channels (n=326), with the mean direction in dark blue. Light blue bars give the mean and black whiskers the standard deviation across electrode bundles (n=42). V-tests assessed non-uniformity around 180 degrees, and were compared against 500 time-shuffled datasets (permutation test); **C**: Event-related potentials (ERPs) for encoding (red) and retrieval (yellow) locked to the patient’s response. Solid lines give the mean, shaded areas the SEM. Grey dots indicate time windows with significant phase opposition between encoding and retrieval (V-test in 200 ms sliding windows spaced 10 ms apart; FDR-corrected with q=0.05). Red and yellow dots are the time points of peak PPC for individual patients used in Figure 6B. ns: not significant; *: 0.01<p<0.05; **: 0.0001<p<0.01.

## 5. Discussion

In this study we demonstrated that oscillations can be detected in behavioral responses from associative memory tasks. Using the Oscillation score, a method to detect oscillations in spike trains (Muresan et al., 2008), we showed that button presses that indicate the timing of encoding and retrieval of a memory were rhythmically modulated, i.e. periodically more or less likely to occur, predominantly in the 1-5 Hz frequency band. We found no evidence for behavioral oscillations for task phases that do not depend on memory. Button presses from forgotten trials did not lock to the oscillation of remembered trials, a distinction that was echoed by hippocampal LFP recordings from 10 epilepsy patients: phase consistency across trials significantly increased in the slow theta range during the encoding and retrieval of correctly, but not incorrectly, memorized objects. Finally, we showed that phase consistency during encoding and retrieval peaked at opposite phases of the theta cycle, aligning with earlier work suggesting that encoding- and retrieval-related information flows are orchestrated by the phase of the hippocampal theta rhythm. Our data extend previous findings by showing that these hippocampal mechanisms influence the timing of overt human behavior.

In our study, we relied on button presses that explicitly marked the timing of memory formation and recall. Though the timing of these button presses is arguably subjective and relies on multiple neural processes, our results allow us to exclude several alternative explanations. Firstly, the behavioral oscillations could not be explained by rhythmicity in visual processing, as the Encoding and Visual task phases shared identical visual inputs. Secondly, a behavioral oscillation was detectable when memory reinstatement was combined with a catch question (Catch-with-retrieval), but not when the catch question was asked 3 seconds after reinstatement (Catch-after-retrieval), suggesting that 1) the observed oscillation did not result from motor processes and 2) the lack of oscillations in memory-independent phases cannot be attributed to the nature or content of the catch questions. The data also show that rhythmic clocking is not universal within memory tasks: while correct trials showed locking to a theta oscillation, incorrect trials did not, a result that was mirrored in electrophysiology.

Our behavioral results, combined with the hippocampal recordings, suggest a strong link between the oscillations in the button presses and the underlying memory processes in hippocampus. We did not observe significant oscillations in behavior for processes that we a priori marked as memory-independent, namely answering the catch questions after reinstatement and the visual task. We did however find increased PPC in hippocampal signals after the catch question appeared on the screen. Interestingly, the O-scores for the corresponding Catch-after-retrieval task phase, although not significant in themselves, were higher than for the visual task. A possible explanation is that retrieval-induced oscillations extend in time, or that catch questions induce a second, weaker reinstatement of the memory, leading to behavioral oscillations that are too weak to detect robustly. Alternatively, the oscillations observed for the catch questions could result from maintaining the retrieved object in working memory. Working memory has been proposed to be mediated by theta-nested gamma bursts (Lisman and Idiart, 1995; Lisman and Jensen, 2013), synchronizing a network of memory areas in frontal, temporal and parietal cortices (Bahramisharif et al., 2018; Fuentemilla et al., 2010; Jacobs et al., 2006; Raghavachari et al., 2001; Vilberg and Rugg, 2012), as well as the hippocampus (Axmacher et al., 2010; for reviews see Hsieh and Ranganath, 2014; Roux and Uhlhaas, 2014). The micro-electrode recordings presented here do not allow us to distinguish between retrieval-related theta oscillations and theta-rhythmic interactions with hippocampus during working memory maintenance. Yet our data do suggest that, at least, retrieval-related theta oscillations are associated with rhythmicity in behavior. Further work is needed to shine light on the extent to which oscillations occur during the catch question phase and to clarify their origin and function.

The behavioral theta oscillations for memory-dependent task phases, together with increased PPC in hippocampal LFPs across trials, suggest that events in the memory task (i.e. cue/stimulus onset) induce consistent phase resets in the hippocampal theta rhythm. Our findings suggest this phase reset is most pronounced in the slow 1-5 Hz theta band. Several human intracranial EEG studies have reported prominent slow theta oscillations during episodic memory tasks (Goyal et al., 2020; Griffiths et al., 2019; Jacobs, 2014; Lega et al., 2012), while the higher 4-8 Hz frequency band typically observed in rodents seems to be linked to movement or spatial processing (Goyal et al., 2020). LFP phase resets and phase-locking after task events have been reported for the slow theta band in memory paradigms (Haque et al., 2015; Mormann et al., 2005; Rizzuto et al., 2006; Rutishauser et al., 2010), and phase consistency directly preceding (Fell et al., 2011) and following (Fell et al., 2008) stimulus presentation has been shown to predict memory performance. Like in rodents, firing of human hippocampal neurons locked to theta oscillations shortly before and during the encoding of later-recognized but not later-forgotten images, for both slow and fast theta bands (Rutishauser et al., 2010). In line with our findings, theta phases were found to differ between encoding and retrieval task phases (Rizzuto et al., 2006), although the effects were limited to an early time window after stimulus presentation, and theta frequencies below 4 Hz were not included in this study. Our intracranial EEG results extend these previous findings by demonstrating that post-stimulus phase consistency extends in time in a narrow frequency band, providing a potential neurophysiological mechanism for the theta-clocked behavior.

Although our behavioral results align closely with the PPC analyses in terms of dominant frequency and subsequent memory effect, our data cannot establish a causal relationship between the hippocampal and behavioral rhythms. Optogenetic and pharmacological techniques allow for such experiments in rodents (McNaughton et al., 2006; Siegle and Wilson, 2014), while patients with hippocampal and/or medial septal lesions are a possible route to establishing causality in humans. Lesion studies can however have profound confounds, such as widespread disruptions in network dynamics beyond the hippocampal theta rhythm and substantial memory deficiency. Interestingly, a new series of studies using transcranial magnetic stimulation over lateral parietal cortex might provide a new way of establishing causality. TMS over lateral parietal cortex was shown to improve memory performance (Kim et al., 2018; Wang et al., 2014), particularly when stimulating in theta-bursts (Hermiller et al., 2019, 2020), possibly due to increased MTL activity and/or increased coherence between hippocampus and the posterior memory network. If theta-frequency TMS is able to enhance memory performance by boosting or entraining the theta rhythm, this approach could potentially establish a direct link between hippocampal and behavioral oscillations in healthy humans.

In recent years, phase coding has appeared as a powerful neural mechanism to optimize specificity and sensitivity on the one hand, and flexibility on the other, and has been shown in several cognitive domains. In addition to sampling of to-be-encoded and retrieved information in hippocampus, rhythmic switching of visual attention has been demonstrated at theta frequencies (Busch and VanRullen, 2010; Fiebelkorn et al., 2018; Helfrich et al., 2018; Landau and Fries, 2012). In memory tasks, potentially interfering mnemonic information has been shown to recur at different phases (Kerren et al., 2018; Kunz et al., 2019; Staudigl and Hanslmayr, 2013) or cycles (Kay et al., 2020) of the hippocampal theta rhythm. Finally, items kept in working memory are thought to be represented in gamma cycles separated in theta/alpha phase (Bahramisharif et al., 2018; Lisman and Jensen, 2013). Visual stimulation at relevant theta/alpha phases, but not opposite phases, boosted working memory performance (Ten Oever et al., 2020). Similarly, when distractors were presented during a working memory task, accuracy fluctuated with the timing of distractor onset, producing a 2.5 Hz oscillation (Wöstmann et al., 2020). Evidence is thus accumulating in both humans and other animals for a powerful role of phase coding in cognitive processes.

Detecting oscillations in sparse behavioral data is not a trivial task, particularly in memory paradigms that rely on one-shot learning, like the task presented here. The trial counts for these tasks are limited by the number of unique trials participants can perform, which ultimately limits the detectability of oscillations. We showed that, despite these limitations, the O-score method (Muresan et al., 2008) and our Z-scoring approach were sensitive enough to reliably detect oscillations where present, while maintaining sufficient selectivity (i.e. producing non-significant O-scores when an oscillation could not be detected reliably), as demonstrated for our simulated datasets (Supplementary Information). It is important to note, however, that a reliable analysis of oscillations in sparse data requires repeated measurements, either in the form of repeated trials (Muresan et al., 2008), or across a large number of participants, like we have done here.

Our results suggest that theta-rhythmicity of memory encoding and retrieval processes can not only be found in neural correlates, but also has a clear behavioral signature: the likelihood that a memory is being formed or recalled rhythmically fluctuates within a trial, at a slow theta frequency, resulting in rhythmicity of button presses relying on these processes. Our findings suggest that behavior can be a relatively straightforward, yet powerful way to assess rhythmicity of neural memory processes, an approach that has the potential to be extended to many other cognitive domains. Together, our behavioral data and hippocampal LFP recordings point to an important mechanistic role for lasting phase consistency in the hippocampal theta rhythm during memory-dependent processing.

## 6. Acknowledgements

This work was funded by starting grant ERC-2016-STG-715714 (STREAM) of the European Research Council to Maria Wimber and consolidator grant ERC-2015-647954 awarded to Simon Hanslmayr. We thank Sophie Watson, Wing Tse, Jonathan Burton-Barr, Emma Sutton, Thomas Faherty, Alexandru- andrei Moise, Laura De Herde, Britanny Lowe, Jessica Davies and James Lloyd-Cox for their help with collecting the behavioral data and Andrew Reid, Gernot Kreiselmeyer and Rüdiger Hopfengärtner for technical support. We are grateful to all participants for donating their time, and in particular thank the patients, their families and the hospital staff, for accommodating our work.

## 7. Author contributions

JLD, JL and MW designed the experiments, all authors were involved in data collection. MtW performed the data analysis and MtW and MW wrote the manuscript. All authors provided feedback on the manuscript.

## 10. Methods

### 10.1. Participants

A total of 216 healthy participants took part in behavioral, EEG and fMRI/EEG studies using the memory tasks described in the next section. A group of 10 epilepsy patients also performed a very similar memory task, more details about this group are given in section ‘iEEG recordings: patients and recording setup’. A separate group of 95 healthy participants completed the visual tasks. All healthy participants volunteered to participate in the studies and were compensated for their time through a cash payment (£6-8 per hour) or the University’s course credit system. All participants gave written informed consent before starting the study. None of the healthy participants reported a history of neurological or psychiatric disorders and all had normal or corrected-to-normal vision. Participants only took part in one version of the task, e.g. participants in the behavioral visual task could not take part in the memory EEG study. Only the behavioral data are presented here. A subset of the behavioral data (visual experiments 1 & 2, and memory experiments 5 & 6), as well as the EEG data from experiment 10 (see Table S1), were previously reported in (Linde-Domingo et al., 2019). All studies with healthy participants took place in facilities of the University of Birmingham, and the participants were recruited through the university’s research participation scheme. All studies were approved by the Science, Technology, Engineering and Mathematics Ethical Review Committee of the University of Birmingham. Demographic information for each of the participant groups is available in Table S1.

### 10.2. Task versions

In this manuscript we present behavioral and intracranial EEG data recorded during a series of visual and memory experiments. The experiments were originally designed to address the following question: is perceptual information about a stimulus analyzed earlier or later than semantic information, and is this processing order similar when viewing a stimulus compared with reinstating the same stimulus from memory? Data from 5 experiments and the analyses addressing the original research question have previously been reported in (Linde-Domingo et al., 2019). In the present manuscript, we analyze the button presses for perceptual and semantic questions together. We also include the behavioral data from an additional 8 follow-up experiments that took place after the collection of the initial datasets.

The experiments can be divided into 3 main categories (Figure 1): memory reaction time experiments; electrophysiology memory experiments; and visual reaction time experiments. All experiments used similar stimulus sets, which we describe below. We then give a general description of each category of experiments, as well as specific differences between experiments within each category. The numbers of participants per task version and their demographic information is given in Figure 1 and Table S1. The characteristics of each of the 13 task versions are summarized in Table S2.

#### Stimulus sets

Across the experiments, 3 different stimulus sets were used, Standard, Shape and Size (Table S2). Each stimulus set consisted of 128 emotionally neutral, everyday objects. Each object fell into one of two perceptual categories and one of two semantic categories. Participants were instructed about the perceptual and semantic categorizations before onset of the study and were shown examples that were not included in the remainder of the study. In the standard stimulus set, used in most experiments, the semantic dimension divided the objects into animate and inanimate objects, while in the shape and size stimulus sets used in experiments 3, 4, 7 and 8, objects were categorized as natural or man-made. Furthermore, 3 different perceptual dimensions were used across the tasks. In the standard stimulus set, half of the stimuli were colored photographs and the other half were black-and-white drawings. In the shape and size stimulus sets only colored photographs were used. Instead, stimuli were categorized as either long or round objects (shape stimulus set, exp. 3 and 7), or stimuli were presented as large or small pictures on the screen (size stimulus set, exp. 4 and 8).

#### Groups 1 & 2: Memory experiments

In the memory experiments, participants first learned associations between cues and objects and later, after a distractor task, memories were reinstated in a cued recall phase, described in more detail below. Participants learned a total of 128 associations divided into blocks of between 4 and 8 trials. Each block consisted of an encoding phase, a distractor phase and a retrieval phase. Cues consisted of action verbs (e.g. spin, decorate, hold, …) for all experiments except experiment 12 (details below).

Each encoding trial started after presentation of a fixation cross for between 500 and 1500 ms to jitter the onset of the trial. The cue then appeared in the center of the screen for 2 s. After presentation of a fixation cross for 0.5-1.5 s the stimulus (stimuli in experiment 6) appeared. Participants were asked to indicate when they made the association between cue and stimulus by pressing a button (encoding button press). The stimulus remained on the screen for 7 s. After the encoding phase, the participants performed a distractor task in which they judged whether numbers presented on the screen were odd or even. The distractor task lasted 60 s. In the retrieval phase the participants were presented with the same cues as during encoding, though in a randomly different order, and asked to recall the associated objects. They then answered either a perceptual or the semantic question about the reinstated object. The trial timed out if the participant did not answer within 10 s. Trials were separated by a fixation cross for 500-1500 ms.

The structure of the retrieval phases differed slightly between experiments. We therefore make a further distinction within the memory experiments, into the electrophysiology experiments (group 1; experiments 10-13) and the behavioral experiments (group 2; experiments 5-9).

For group 1, we aimed to separate the reinstatement processes from the formulation of the answer. To this end, participants were asked to indicate, through a button press, when they had a clear image of the associated object in mind. The trial timed out if the participant did not press the button within 10 s. They then kept the image in mind for 3 seconds. Finally, the question and answer options appeared on the screen, after which the participants responded as quickly as possible. Participants had 3 s to respond. As a result, the retrieval trials of the electrophysiology experiments produced two button presses: a retrieval button press and a catch-after-retrieval button press. These button presses are analyzed separately. Only the reinstatement button press is considered memory-dependent, because the catch question appears at a time point when the object has supposedly already been fully retrieved.

For group 2, the answer options were shown on the screen for 3 s before the retrieval cue appeared. The catch-with-retrieval button presses obtained for the memory reaction time experiments can therefore be assumed to represent the time point when sufficient information has been retrieved about the object to answer the catch question.

The number of times we asked participants to retrieve associations was varied between behavioral experiments in group 2. In experiments 5, 7 and 8, every object was probed twice, and participants answered both the perceptual and semantic question for each object in random order. In experiment 9, every object was reinstated 6 times in the retrieval phase of the block, and twice during a delayed retest 2 days later, with the delayed test not included due to poor performance (average performance 49.6%, only 16 out of 52 participants performed above chance). In experiment 6, participants learned associations of triplets instead of pairs, consisting of cue, object and scene image. During the retrieval phase of this experiment, each object was probed only once, and in addition to the perceptual and semantic questions participants were asked a question about the background image (indoor or outdoor?), such that each question was answered on 1/3 of the trials.

The group 1 memory task was used for EEG recordings (experiment 10), combined EEG/fMRI recordings (experiment 11) and intracranial EEG recordings in epilepsy patients (experiments 12 and 13, also see section 10.3). Several small adjustments were made to the task to accommodate electrophysiology. First, to minimize the duration of the testing sessions, the duration of the distractor phase was reduced to 20 s in the EEG/fMRI experiment, while in the EEG and iEEG task versions, every object was reinstated only once. To compensate for the corresponding drop in the number of catch questions, participants answered both perceptual and semantic catch questions on every trial, one after the other, in random order. The doubling of the number of catch questions per trial was introduced after the first 3 EEG participants and 3 iEEG patients were recorded. In addition, the first 3 iEEG patients learned pairs of background scenes and objects, instead of verb-object pairs, with the background scenes functioning as cues during the retrieval phase (experiment 12). These patients only learned a total of 64 pairs. The reaction time data from these 3 patients showed no difference to that of the other 7 patients (see Figure S1) and no qualitative differences were found in the PPC analysis (PPC per patient is shown in Figure S4). During encoding trials, the background and object appeared on the screen at the same time. As a result, the encoding trials of these 3 participants do not have a separate cue period.

We made two further modifications to the task for the iEEG recordings in all epilepsy patients. First, the task was made fully self-paced, such that length of verb presentation and the period needed to associate cue and object were determined by the patient on each trial. The patients pressed a button when they were ready to move on. Second, to avoid loss of attention/motivation and/or to accommodate medical procedures, visitors and rest periods, the task was divided into two or three sessions, recorded at different times or on different days. Data from different sessions were pooled and analyzed together. Details of the electrophysiological recordings included in this manuscript can be found in the section ‘iEEG recordings: patients and recording setup’.

#### Group 3: Visual reaction time experiments

In the visual experiments, participants were shown a series of stimuli on the screen, and were asked either a perceptual or a semantic question about each stimulus. The stimuli and questions used in the visual experiments were identical to those in the memory experiments. To get accurate estimates of the reaction times, the answer options were shown for 3 s prior to stimulus presentation. Stimuli were presented in the center of the screen. Each trial was preceded by a fixation cross for a random duration of between 500 and 1500 ms, so the onset of the trial could not be predicted. Like in memory group 2, participants were instructed to answer as fast as possible.

In experiments 1, 3 and 4, all 128 stimuli were shown twice, once followed by a perceptual and once by a semantic question (in random order), so both questions were answered for every object. In experiment 2, in which the object images were shown with a background, all stimuli were presented only once, followed by one of three questions: perceptual, semantic or contextual, with the later referring to the background (indoor or outdoor). All button presses were included here. We refer to (Linde-Domingo et al., 2019) for analyses comparing the different catch questions.

### 10.3. Assessment of performance and exclusion of participants

Prior to reaction time analyses, the performance of each of the participants was analyzed based on their accuracy in answering the catch questions. Answers to catch question were considered incorrect when subjects chose the wrong answer, when the indicated they had forgotten the answer (for memory tasks) or when they did not answer on time (for healthy participants only). The data of a participant were only included in the analysis of a task phase if the following two requirements were met: 1) catch question accuracy across trials exceeding chance level and 2) a minimum of 10 correct button presses per participant in the task phase of interest. The first criterium was assessed using a one-sided binomial test against a guessing rate of 50% with α = 0.05. The second criterium had to be set as some participants repeatedly failed to provide encoding (reinstatement) button presses before trial time-out, leaving too few trials to run further analyses for the Encoding (Retrieval) phase, despite sufficient performance when answering the catch questions. The inclusion criteria were set *a priori*. The number of participants included in each of the task phases is shown in Figure 1 and the number of excluded participants can be found in Figure S1A.

Of the participants who performed the memory tasks, 28 participants answered two catch questions per retrieval trial, while the remaining 198 answered one. To bring the analyses of the 28 participants with 2 catch questions per trial in line with the data from the other 198 participants, we considered a 2-catch trial to be correctly reinstated if one or both catch questions were answered correctly (on average, across 28 subjects: one catch question correct: 12.5% of trials; two catch questions correct: 76.0% of trials, see Figure S1D).

### 10.4. RT analysis: O-score and statistics per participant

To assess the presence and strength of oscillations in behavioral responses we used the Oscillation score (O-score, Figure 2B), a method that was developed to analyze oscillations in spike trains (Muresan et al., 2008). Like spikes, the button presses we study here are discreet, all or nothing events, and can be summarized as trains of button presses across trials. The O-score method identifies the dominant frequency in those trains, and produces a normalized measure of the strength of the oscillation that can be compared across conditions.

We added an additional processing step before computing the O-score, to compensate for the fact that behavioral responses, unlike spikes, have no baseline rate (e.g. they cannot occur before cue/stimulus onset). Extremely early and late responses therefore have to be considered outliers. We removed these outliers prior to O-score computation by removing the first and last 5% of the button press trace of each participant, i.e. maintaining the middle 90% of the button presses. Figure S2B shows that reducing the fraction of button presses included in the analyses affected the ability to identify oscillations, but did not affect the differences found between the task phases.

The button presses from correctly answered trials that remained after outlier removal entered the O-score computation. We made two modifications to the procedure described in (Muresan et al., 2008) to match the characteristics of our dataset. The O-score procedure and our modifications are described below.

We defined a wide frequency range of interest of between 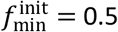 and 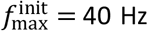 for the O-score analysis. Given the wide variety in the number of responses and response times, we checked for every participant and task phase whether these pre-set frequency bounds were valid. Following (Muresan et al., 2008), we increased the lower bound to 1/*c_min_* of the width of the response distribution (in seconds) of the participant, with *C*_min_ = 3, such that at least 3 cycles of the lowest detectable frequency were present in the data. We reduced the upper bound to the average response rate (button presses per second), if the participant did not have enough button presses to resolve the upper frequency limit. The O-score was then computed through the following series of steps:

Step 1: As described in (Muresan et al., 2008), we computed the auto-correlation histogram (ACH) of the button presses with a time bin size of 1 ms (*f*_*S*_ = 1000 Hz).

Step 2: The ACH was smoothed with a Gaussian kernel with a standard deviation *σ*_fast_ of 2 ms. As estimated in (Muresan et al., 2008), this smoothing kernel attenuated frequencies up to 67 Hz by less than 3 dB, allowing us to detect frequencies in the entire frequency range of interest.

Step 3: We identified the width of the peak in the ACH using the method described in (Muresan et al., 2008). However, to avoid the introduction of low frequencies by replacing the peak, we opted to only use positive lags beyond the detected ACH peak for further steps, as the peak-replacement approach would not allow us to detect frequencies toward the lower bound of our frequency range of interest. To identify the peak, we smoothed the ACH with a Gaussian kernel with *σ*_slow_ of 8 ms, resulting in the smoothed ACH trace *A*_slow_(*l*), with *l* the lag. We then identified the left boundary lag of the central peak *l*_left_ by:

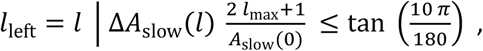

Where *l*_max_ is the highest lag included in the ACH and *A*_slow_(0) is the value of the peak of the ACH (i.e. at lag 0).

Step 4: The remaining part of the ACH was subsequently truncated/zero padded to size *w*, where 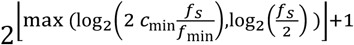. We then applied a Hanning taper and the Fourier transform was computed.

Step 5: We identified the frequency with the highest power in the participant-adjusted frequency bounds, as well as the average magnitude of the spectrum between 0 and *f*_*S*_/2 Hz. The O-score was then computed as: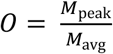.

In their paper, Muresan and colleagues propose a method to estimate the confidence interval of the O-score, allowing for a statistical assessment at the single cell level. However, this approach requires multiple repeated recordings, which are not available for the data presented here, nor do the datasets contain enough data points to create independent folds. Instead, we opted to generate a participant-specific reference distribution of O-scores for the identified frequency, to which we could compare the observed O-score. To this end, we randomly generated 500 time series for each participant matching the trial count and overall response density function of the participant’s original button presses. First, a gamma probability function *r*_gamma_(*t*) was fitted to the participant’s response distribution and scaled to the number of responses of the participant. We then generated 500 Poisson time series, with the probability of a response in a time step Δ*t* = 0.5 ms, given by:

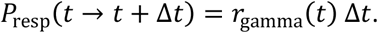

If a gamma distribution could not be fitted (as assessed through a *χ*^2^ goodness-of-fit test with α = 0.05), the participant’s button presses were instead randomly redistributed in time, with the new time per button press uniformly drawn from a window defined by the participant’s peak frequency and centered around the original response time.

O-scores were then computed for each of the resulting reference traces, but instead of finding the peak, the power at the peak frequency of the observed O-score was used. This approach controls for any frequency bias that could arise due to the length of the time series and/or the number of datapoints included in the analysis. To compare the observed O-score to the reference O-scores, we first log-transform all O-score values. This log-transformation was needed as the O-score is a bounded measure (it cannot take values below 0) and the O-score distribution is therefore right-skewed when O-score values are low, leading to an underestimation of the standard deviation of the reference distribution. The log-transformed reference O-scores were then used to perform a one-tailed Z-test for the observed O-score at α = 0.05, establishing the significance of the oscillation at the single participant level. For a validation that 500 reference O-scores was sufficient to produce a stable outcome for the Z-scoring, we refer to Figure S2A. Second level t-scores were subsequently computed based on the Z-scored O-scores for each task phase and tested with α = 0.01 (one-tailed, Bonferroni-corrected for 5 task phases). These Z-scored oscillation scores can be assumed to represent the strength of the behavioral oscillation, and are the basis of many of our statistical comparisons.

To test whether the O-scores of memory-dependent task phases, i.e. Encoding, Retrieval and Catch-with-retrieval, and memory-independent task phases, i.e. Catch-after-retrieval and Visual, against each other, we fitted a linear mixed model to the Z-scored O-scores, with memory-dependence and the length of the time series used for O-score computation as fixed effects, and an intercept per participant as random effect. We included the length of the time series, computed as the difference (in seconds) between the last and the first RT used in the O-score analysis, because there was a substantial difference in response times between the task phases, with overall similar patterns as the O-scores (see Figure S1). We included participants as random effects to compensate for the difference in the number of datapoints contributed by memory task participants (3 datapoints from group 1, 2 datapoints from group 2) compared to visual task participants (1 datapoint from group 3), and to account for dependencies in the data. We fitted an identical linear mixed model to the peak frequencies corresponding to significant O-scores. The linear mixed models were fitted using the fitlme function from the Statistics and Machine Learning Toolbox for MATLAB (The Mathworks Inc.).

The performance of the modified O-score method and Z-scoring procedure were tested in a simulated dataset where the amplitude and frequency of the oscillation in the simulated button presses was varied. Methods and results of these simulations are given in the Supplementary Information.

### 10.5. RT analysis: Phase of response

For the task phases with significant second level O-scores, i.e. Encoding, Retrieval and Catch-with-retrieval, we analyzed the phases at which individual button presses occurred in the behavioral oscillation identified by the O-score analysis. We performed this analysis for both correctly and incorrectly remembered trials. As this analysis relied on the frequency identified by the O-score analysis, only participants with significant O-scores were included.

To identify the phases of the button presses, we first established a continuous reference trace that captured the behavioral oscillation. This was achieved by convolving the button presses with a Gaussian kernel, with *σ*_freq_ = *f*_peak_/_8_. The resulting continuous trace was then band-pass filtered with 2^nd^ order Butterworth filter with a 1 Hz wide pass band centered on the participant’s peak frequency identified by the O-score. The filtered trace was then Hilbert transformed and the instantaneous phase was computed, resulting in a phase of 0 rad for the peak of the behavioral oscillation. Finally, for each button press, the corresponding phase of the reference trace was determined and stored for further analyses.

We used two complementary approaches to compare the phase-locking of correct versus incorrect trials: across participants, allowing us to include correct and incorrect trials from all participants with significant O-scores, even when the number of incorrect button presses was low; and within participants, comparing the phase distributions of correct and incorrect trials for participants with 10 or more incorrect trials. These approaches are described in more detail below.

With the across-participant analysis we aimed to address the following questions: 1) are correct and incorrect trials phase-locked to the behavioral oscillation found for the correct trials and 2) are correct trials locked to this oscillation more strongly than incorrect trials? For these analyses, to find the phases of incorrect trials, we compared the timing of the incorrect button presses to the phase trace determined on the correct trials only. To determine the phases of the correct trials, to avoid circularity, we instead used a leave-one-out approach; for each correct button press, a phase trace was established based on all other correct trials. We then performed a V-test (implementation: CircStat toolbox; Berens, 2009) to assess non-uniformity of the phase distributions around the peak of the behavioral oscillation (i.e. around phase 0 rad), providing an answer to the first question. To address the second question, i.e. whether correct phase distributions were modulated more strongly than incorrect phase distributions, we had to compensate for the trial count differences as well as the methodological differences in determining the phase distributions for correct and incorrect trials. To this end, we defined the permutation test statistic *V*_diff_ = *V*_correct_ − *V*_incorrect_, with *V* being the test statistic from the V-test for non-uniformity around phase 0. For each participant with a significant O-score, we then randomly shuffled the labels of the correct and incorrect trials, and computed the *V*_diff_ statistic across participants for the label-shuffled trials in the same way as described for the observed labels. We repeated this shuffling procedure 100 times and counted the number of times 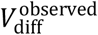 was smaller than 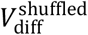. This procedure hence resulted in a p-value that estimated the likelihood that the observed difference in phase modulation between correct and incorrect trials was produced by chance.

For participants with sufficient (10 or more) incorrect trials, we performed an additional analysis to compare the phase modulation of correct and incorrect trials. For these participants, the correct trials were randomly subsampled to match the number of incorrect trials. The phases of the incorrect trials and the subsampled correct trials were then determined based on the phase trace of the remaining correct trials and V-tests were performed for both subsampled correct and incorrect phase distributions. The V-statistics for correct trials were then compared to those for incorrect trials using paired t-tests. The subsampling procedure was repeated 100 times.

### 10.6. iEEG recordings: patients and recording setup

We recorded intracranial EEG from 10 epilepsy patients while they were admitted to hospital for assessment for focus resection surgery; 7 patients were recorded in the Queen Elizabeth Hospital Birmingham (Birmingham, UK) and 3 patients in the Universitätsklinikum Erlangen (Erlangen, Germany). For an 11^th^ patient, task recording was aborted due to poor performance. All patients were recruited by the clinical team, were informed about the study and gave written informed consent before their stay in hospital. Ethical approval was granted by the National Health Service Health Research Authority (15/WM/2019), the Research Governance & Ethics Committee from the University of Birmingham, and the Ethik-Kommission der Friedrich-Alexander Universität Erlangen-Nürnberg (142_12 B).

The patients were implanted with between 2 and 8 Behnke-Fried electrodes with microwire bundles (AdTech Medical Instrument Corporation, USA) in the medial temporal lobe (see Figure 1 for electrode placement and Table S6 for electrode numbers per patient), as well as a number of depth electrodes in other brain areas. Only data from the microwires are presented here, since these were targeted at hippocampal grey matter. Implantation schemes were determined by the clinical team and were based solely on clinical requirements. Each microwire bundle contained 8 high-impedance wires and 1 low impedance wire, which was used as reference in most patients (see Table S6 for patient-specific references). Data were recorded using an ATLAS recording setup (Neuralynx Inc, USA.) consisting of CHET-10-A pre-amplifiers and a Digital Lynx NX amplifier. Data were filtered using analog filters with cut-off frequencies at 0.1 Hz and 9000 Hz (40 Hz for patient 01) and sampled at 32,000 Hz in Birmingham and 32,768 Hz in Erlangen. All data were stored on the CaStLeS storage facility of the University of Birmingham (Thompson et al., 2019).

For each patient both pre- and post-surgical T1-weighted MRI images were acquired. The pre- and post-surgical scans were co-registered and normalized to MNI space using SPM12. The locations of the tip of the macro-electrodes were determined through visual inspection using MRIcron and electrodes were assigned one of the following anatomical labels: amygdala, anterior, middle or posterior hippocampus, or parahippocampal gyrus. The locations and labels were visualized using ModelGUI and are shown in Figure 5A.

The patients performed the memory task described in section ‘Task versions’ on a laptop computer (Toshiba Tecra W50), while seated in their hospital bed or on a chair next to their bed. The three patients who were recorded in Erlangen, Germany, performed the task in German. Patients completed between 64 and 128 full trials, divided over between 1 and 3 recording sessions (see Table S7). Of the 10 patients, 3 patients performed a version of the task that used scene images as cue (see above), while the other patients were presented with verbs as cues. For the image cue task version, the cue was shown at the same time as the object, hence the encoding data of 3 patients had no separate cue phase.

### 10.7. iEEG analysis: LFP data preprocessing

Raw microwire data were loaded into MATLAB using the MatlabImportExport scripts (version 6.0.0) provided by Neuralynx Inc.. The data were subsequently zero-phase filtered with a 3^rd^ order FIR high-pass filter with a cut-off frequency of 0.5 Hz and a 6^th^ order FIR low-pass filter with a cut-off frequency of 200 Hz using FieldTrip (Oostenveld et al., 2011). A Notch filter with a stopband of 0.5 Hz wide at −3 dB was used to remove 50 Hz line noise and its harmonics. The data were down-sampled to 1000 Hz and divided into encoding and retrieval trials.

All data were visually inspected and channels/time points that contained electrical artefacts or epileptic activity were removed. Trials that had more than 20% of time points marked as artefactual were rejected in their entirety. In an additional pre-processing step, the data of patient 03 were re-referenced against the mean of the channels in each microwire bundle. This was done to bring the data from this patient, whose data were originally recorded against ground, more in line with the referencing schemes of the other patients, which were recorded against a local reference wire (see Table S6 for reference information per patient).

### 10.8. iEEG analysis: Wavelet transform, Pairwise Phase Consistency and Cluster statistics

The pre-processed microwire recordings were wavelet transformed using a complex Morlet wavelet with a bandwidth parameter of 4. We used the cwt implementation from the Wavelet Toolbox for MATLAB (The Mathworks Inc., USA) to compute the wavelet transform. The wavelet was scaled to cover a frequency range between 1 and 12 Hz in 43 pseudo-logarithmic steps and convolved with the data in time steps of 10 ms.

To obtain the power plots in Figure S4C, we extracted the absolute value of the wavelet coefficients and assessed power changes per frequency against a −2 to −0.5 s pre-cue baseline using a two-sided t-test for every time point. We averaged the resulting t-maps across the wires within each bundle, as they shared a common low impedance reference. The bundle averages were then used to compute a second level t-score across the bundles of all participants. The p-values resulting from the second level analysis were entered into a Benjamini-Hochberg false discovery rate (FDR) correction procedure with q = 0.05 to correct for multiple comparisons and the t-score map was masked at alpha = 0.05.

The phases obtained for every frequency and time point in the trial using the complex wavelet transform were used to compute the pairwise phase consistency (PPC, Vinck et al., 2010) across trials for each time- and frequency pixel and for each microwire. The PPC was calculated for correct and incorrect trials separately. The PPC values were then non-parametrically tested relative to their pre-cue baseline, defined as the period from 2 s to 0.5 s prior to cue onset, using a Mann-Whitney U-test. We opted for a non-parametric test due to the strong left-skew of the PPC data. As for the power analyses, the resulting approximated Z-values were averaged across the microwires in a bundle, and the averages were used to compute a second level t-score across all bundles from all patients.

We then detected time-frequency clusters of significant PPC through the following steps. First, the t-scored PPC values were thresholded at α = 0.05 with df = *N*_bundles_ − 1, resulting in a binary image with 0 = non-significant and 1 = significant. This binary image was entered into an 8-connected component labelling algorithm to identify clusters of significant PPC values.

As we used a fixed threshold to identify the clusters, it is possible for clusters to be made up of two or more merged peaks. This merging of peaks artificially inflates the cluster’s size. To avoid this, we tested whether each cluster contained more than one peak, and if so, split the cluster. To this end, for every cluster, we iteratively increased the significance threshold towards 90% of the highest value in the cluster, in 5% increments, and reran the cluster detection method described in the previous paragraph. We required any resulting subclusters to be at least 5% of the size of the original cluster, to overcome noise in the data. If no subclusters were found, the threshold was increased further. On the other hand, if all identified subclusters were smaller than 5% of the original cluster, we concluded that the cluster could not be split. If subclusters of sufficient size were detected, these were stored. For all pixels that were part of the original cluster, but were not a member of any of the new subclusters, we computed the weighted Euclidian distance to all subclusters and assigned them to the closest subcluster. For each resulting (sub)cluster we then computed a cluster statistic defined as the sum of all t-scores from all pixels in the cluster.

We took a non-parametric approach to assess the statistics at the cluster level. To this end, we went back to the wavelet transforms and, at a random time point in each trial, divided the trial in two parts. We then concatenated the first part of the trial to the end of the second part. This procedure, suggested in (Cohen, 2014), left all characteristics of the dataset intact, with the exception of the temporal structure of the phase. We computed the PPC across these time-shuffled trials, Z-scored against baseline, computed the second level t-score, identified clusters of significant t-scores and computed the cluster scores as described in the previous paragraph. We repeated this procedure 100 times and we stored the highest cluster score for each repetition, resulting in a reference distribution of maximum cluster scores. We then non-parametrically compared the cluster scores from the intact data to the reference distribution, with α = 0.05. We performed the time-shuffle analysis independently for positive and negative changes in PPC and for correct and incorrect trials separately.

Finally, we also compared the PPCs from correct and incorrect trials to each other directly. We used a similar approach as described above, with two important differences: 1) the second level analysis was now performed on the pair-wise difference between correct and incorrect PPCs from the same bundle and 2) we shuffled correct and incorrect trials (as opposed to time points) to obtain the reference distribution.

### 10.9. Phase differences and Event-Related Potentials

We used two different approaches to assess whether phases between encoding and retrieval trials differed. First, we tested whether encoding and retrieval phases differed at the moment of peak PPC, where the effect of phase resets was optimal and trials were most phase aligned. To this end, we detected the highest PPC value for every participant and stored the average phase for every electrode at the corresponding time and frequency. We then computed the phase difference per electrode by subtracting the retrieval phases from the encoding phases. This procedure was performed on both the cue- (for retrieval) or stimulus- (for encoding) locked data and for the response-locked data. We subsequently performed V-tests for non-uniformity around 180 degrees to assess whether the phases of encoding and retrieval were opposite at peak PPC. We compared the V-statistics to V-statistics computed using the same approach in 500 time-shuffled datasets (see previous section).

For the second approach we computed Event-Related Potentials (ERPs) to test for phase opposition in time windows leading up to the response. To obtain the ERPs, we first Z-scored the raw data per electrode by subtracting the mean and dividing by the standard deviation of all trials and time points. We then tested whether all electrodes had the same sign. This step was essential because recordings from different layers of the hippocampus can have opposing polarities. In micro-wire recordings there is no control over the placement of the electrode, nor is it possible to determine this placement based on scans, hence potential sign flips have to be detected in the data, before averaging data of different electrodes. We detected the sign by identifying the highest deflection in the trial-average of every electrode in the 1 second time interval after cue onset during encoding. If this deflection was negative, the data from the electrode was flipped. Note that the time interval we used for sign testing was not included in the ERP analysis in Figure 6. We then averaged the trials of all wires within a micro-wire bundle, separating correct and incorrect trials. The data resulting from this step are represented in Figure S4D. We then filtered the averaged data in the theta-frequency band (1-5 Hz; data in Figure 6C) and identified the instantaneous phase using the Hilbert transform. We subtracted the instantaneous phases from the retrieval trials of the phases from the encoding trials for each bundle yielding the instantaneous phase difference. The phase differences were collected in windows of 200 ms (i.e. 1 cycle at the 5 Hz upper bound of the theta band) spaced 10 ms apart and we tested whether the phase differences in each window were non-uniformly distributed around 180 degrees using a V-test. We used a Benjamini-Hochberg False Discovery Rate correction procedure (Benjamini and Hochberg, 1995) with q = 0.05 to account for repeated tests across the time windows.

### 10.10. Software and code

The task presentation and all data analyses were performed using MATLAB 2015a-2018a (The Mathworks), using functions from the Wavelet Toolbox and the Statistics and Machine Learning Toolbox. The following third-party functions and software tools were used:

- Psychophysics Toolbox Version 3 (Releases between January 2017 and April 2019): https://github.com/Psychtoolbox-3/Psychtoolbox-3
- CircStats toolbox 2012a (Berens, 2009): https://github.com/circstat/circstat-matlab
- FieldTrip v20190615 (Oostenveld et al., 2011): https://github.com/fieldtrip/fieldtrip;
- NeuralynxImportExport v6.0.0: https://neuralynx.com/software/category/matlab-netcom-utilities
- SPM12: https://www.fil.ion.ucl.ac.uk/spm/
- MRIcron: https://people.cas.sc.edu/rorden/mricron/index.html
- ModelGUI: http://www.modelgui.org

### 10.11. Data and code availability

Custom functions and scripts used to produce the results presented here are available via https://github.com/marijeterwal/behavioral-oscillations. All behavioral data underlying the results in this manuscript have been made available and from the iEEG dataset, the PPC-values for correct and incorrect trials, including the time- and trial-shuffled PPCs have been made public (ter Wal et al., 2020, https://doi.org/10.6084/m9.figshare.c.5192567). Other data will be made available upon reasonable request, but note that we cannot provide raw iEEG data and patient-specific electrode locations due to privacy and consent restrictions.

## Supplementary Information

### 1. Supplementary Tables S1 – S7

**Table S1.**
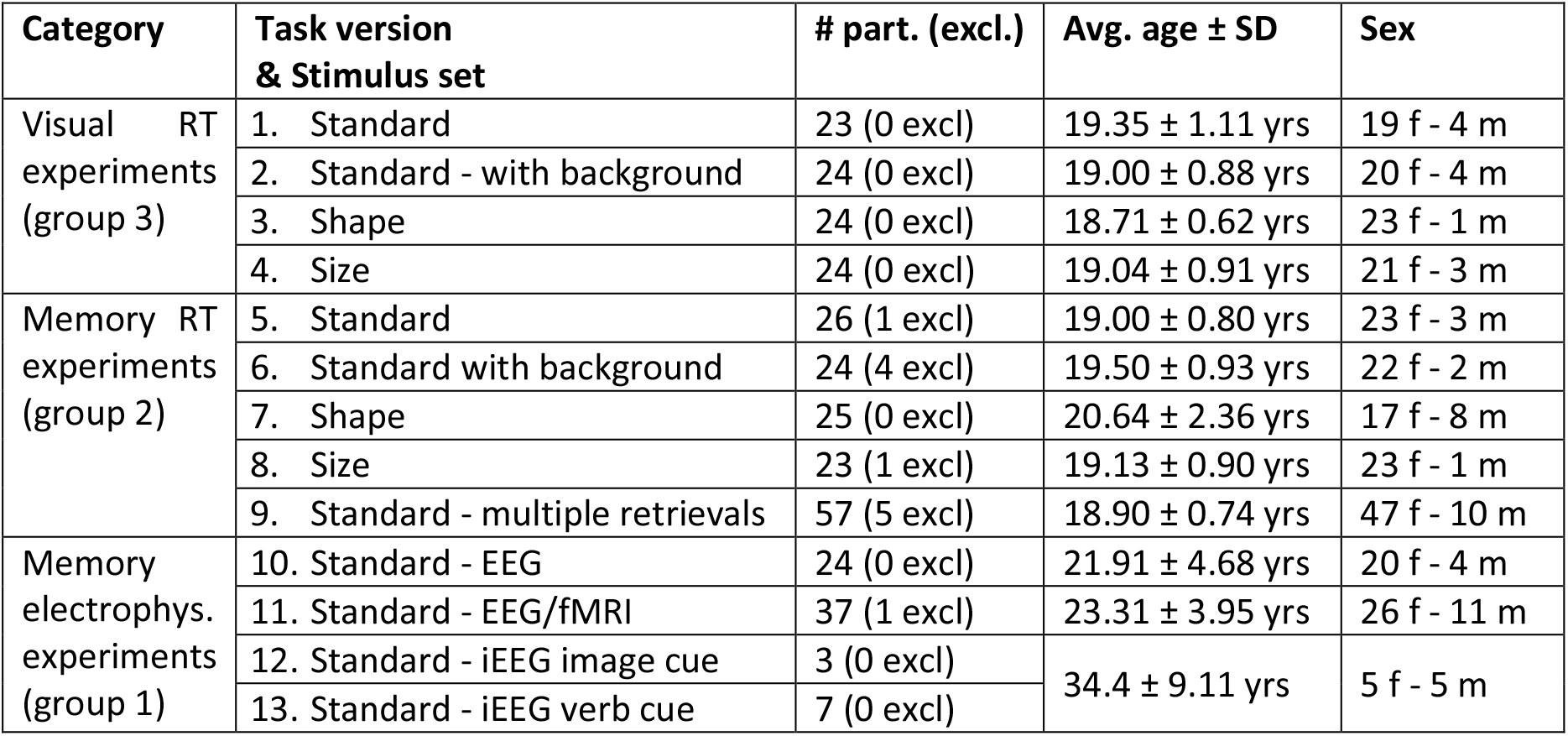
Demographic information for the 13 experiments included in this study. From left to right, the columns provide the following information: category = the general group the experiment fell under (refer to the main Methods section for more details); task version = number of the experiment used throughout the manuscript and stimulus set; # participant (excl.) = the number of participants that completed the experiment, with the number in brackets indicating the number of participants that were excluded due to poor performance for the catch questions; Avg. age ± SD = the mean age and standard deviation of all participants in years; sex = # number of males (m) and females (f) taking part in the study.

| Category | Task version<br>& Stimulus set | # part. (excl.) | Avg. age ± SD | Sex |
| --- | --- | --- | --- | --- |
| Visual RT<br>experiments<br>(group 3) | 1. Standard | 23 (0 excl) | 19.35 ± 1.11 yrs | 19 f - 4 m |
| | 2. Standard - with background | 24 (0 excl) | 19.00 ± 0.88 yrs | 20 f - 4 m |
| | 3. Shape | 24 (0 excl) | 18.71 ± 0.62 yrs | 23 f - 1 m |
| | 4. Size | 24 (0 excl) | 19.04 ± 0.91 yrs | 21 f - 3 m |
| Memory RT<br>experiments<br>(group 2) | 5. Standard | 26 (1 excl) | 19.00 ± 0.80 yrs | 23 f - 3 m |
| | 6. Standard with background | 24 (4 excl) | 19.50 ± 0.93 yrs | 22 f - 2 m |
| | 7. Shape | 25 (0 excl) | 20.64 ± 2.36 yrs | 17 f - 8 m |
| | 8. Size | 23 (1 excl) | 19.13 ± 0.90 yrs | 23 f - 1 m |
| | 9. Standard - multiple retrievals | 57 (5 excl) | 18.90 ± 0.74 yrs | 47 f - 10 m |
| Memory<br>electrophys.<br>experiments<br>(group 1) | 10. Standard - EEG | 24 (0 excl) | 21.91 ± 4.68 yrs | 20 f - 4 m |
| | 11. Standard - EEG/fMRI | 37 (1 excl) | 23.31 ± 3.95 yrs | 26 f - 11 m |
| | 12. Standard - iEEG image cue | 3 (0 excl) | 34.4 ± 9.11 yrs | 5 f - 5 m |
|  | 13. Standard - iEEG verb cue | 7 (0 excl) |  |  |

**Table S2.** Task details for the different experiments. From left to right, the columns provide the following information: task # = number of the experiment used throughout the manuscript; type = experiment type (‘beh.’ for behavioral only); task phases = the task phases this experiment contributed to; Stimulus set = the stimulus set that was used; # trials = the number of unique objects (for visual tasks) or cue-object pairs (for memory tasks) the participants were presented with. For some of the visual tasks, the objects were shown twice, indicated by a ‘x 2’; # retrieval repetitions (for memory tasks only; NA means not applicable) = the number of times each learned object had to be reinstated during the retrieval phases of the experiment; Catch questions: catch questions used for the experiment, with perc. = perceptual, sem. = semantic and cont. = contextual; # catch Q per trial = the number of catch questions that were asked after reinstatement.

| Task # | Type | Task phases | Stimulus set | # trials | # ret. reps. | Catch questions: | # catch Q per trial |
| --- | --- | --- | --- | --- | --- | --- | --- |
| 1 | Beh. | Visual | Standard | 128 x 2 | NA | Perc.: drawing/photo<br>Sem.: animate/inanimate | 1 |
| 2 | Beh. | Visual | Standard background | 128 | NA | Perc.: drawing/photo<br>Sem.: animate/inanimate<br>Cont.: indoor/outdoor | 1 |
| 3 | Beh. | Visual | Shape | 128 x 2 | NA | Perc.: round/elongated<br>Sem.: natural/manmade | 1 |
| 4 | Beh. | Visual | Size | 128 x 2 | NA | Perc.: big/small<br>Sem.: natural/manmade | 1 |
| 5 | Beh. | Encoding<br>Catch-with-ret. | Standard | 128 | 2 | Perc.: drawing/photo<br>Sem.: animate/inanimate | 1 |
| 6 | Beh. | Encoding<br>Catch-with-ret. | Standard background | 128 | 1 | Perc.: drawing/photo<br>Sem.: animate/inanimate<br>Cont.: indoor/outdoor | 1 |
| 7 | Beh. | Encoding<br>Catch-with-ret. | Shape | 128 | 2 | Perc.: round/elongated<br>Sem.: natural/manmade | 1 |
| 8 | Beh. | Encoding<br>Catch-with-ret. | Size | 128 | 2 | Perc.: big/small<br>Sem.: natural/manmade | 1 |
| 9 | Beh. | Encoding<br>Catch-with-ret. | Standard | 128 | 6<br>(7<sup>th</sup> & 8<sup>th</sup> excluded) | Perc.: drawing/photo<br>Sem.: animate/inanimate | 1 |
| 10 | EEG | Encoding<br>Retrieval<br>Catch-after-ret. | Standard | 128 | 1 | Perc.: drawing/photo<br>Sem.: animate/inanimate | 2<br>(1 for 3 part.) |
| 11 | EEG<br>fMRI | Encoding<br>Retrieval<br>Catch-after-ret. | Standard | 128 | 2 | Perc.: drawing/photo<br>Sem.: animate/inanimate | 1 |
| 12 | iEEG | Encoding<br>Retrieval<br>Catch-after-ret. | Standard -<br>image cue | 64 | 1 | Perc.: drawing/photo<br>Sem.: animate/inanimate | 1 |
| 13 | iEEG | Encoding<br>Retrieval<br>Catch-after-ret | Standard -<br>verb cue | 128 | 1 | Perc.: drawing/photo<br>Sem.: animate/inanimate | 2 |

**Table S3.** Reaction time descriptors for correct trials per task phase, across all included participants.

| Measure | Task phase |  |  |  |  |
| --- | --- | --- | --- | --- | --- |
|  | Encoding | Retrieval | Catch-with-retrieval | Catch-after-retrieval | Visual |
| Mean RT (s) | 3.99 | 2.20 | 2.24 | 1.42 | 0.89 |
| SD of RT (s) | 2.57 | 2.22 | 1.26 | 1.25 | 0.45 |
| 5% bound (s) | 1.29 | 0.68 | 0.97 | 0.53 | 0.44 |
| 95% bound (s) | 6.81 | 5.15 | 4.79 | 3.22 | 1.67 |
| Mean # responses per participant | 66.27 | 150.68 | 217.66 | 158.17 | 215.20 |
| SD # responses per participant | 33.93 | 54.28 | 99.2 | 57.88 | 54.1 |

**Table S4.** Effect sizes of the O-scores per task phase (i.e., against the permuted baseline), and between task phases (i.e., comparing against each other).

|  | d' | Cohen's d |  |  |  |  |
| --- | --- | --- | --- | --- | --- | --- |
|  |  | Encoding | Retrieval | Catch-with-ret. | Catch-after-ret. | Visual |
| Encoding | 0.46 |  | -0.21 | -0.12 | 0.23 | 0.92 |
| Retrieval | 0.56 |  |  | 0.064 | 0.39 | 0.92 |
| Catch-with-retrieval | 0.43 |  |  |  | 0.28 | 0.86 |
| Catch-after-retrieval | 0.20 |  |  |  |  | 0.64 |
| Visual | -0.42 |  |  |  |  |  |

**Table S5.** Post-hoc tests for peak frequency from the Oscillation score procedure.

| Group(s) | Comparison | Test (all two-tailed) | Bonferroni-correction | Statistic | p-value |
| --- | --- | --- | --- | --- | --- |
| Group 1 | Encoding vs Retrieval | Wilcoxon signed rank | 3 | 1.081 | 0.839 |
|  | Encoding vs Catch-after-retrieval | Wilcoxon signed rank | 3 | -1.197 | 0.694 |
|  | Retrieval vs Catch-after-retrieval | Wilcoxon signed rank | 3 | -1.776 | 0.227 |
| Group 2 | Encoding vs Catch-with-retrieval | Wilcoxon signed rank | NA | -3.201 | 0.0014 |
| Group 1 & Group 2 vs Group 3 | Encoding vs Visual | Wilcoxon rank-sum | 4 | -4.529 | < 0.0001 |
|  | Retrieval vs Visual | Wilcoxon rank-sum | 4 | -3.664 | 0.000988 |
|  | Catch-with-retrieval vs Visual | Wilcoxon rank-sum | 4 | -2.593 | 0.038 |
|  | Catch-after-retrieval vs Visual | Wilcoxon rank-sum | 4 | -1.340 | 0.721 |

**Table S6.** Implantation information per patient: the number of Behnke-Fried microwire bundles that were implanted in hippocampus (in brackets the number of bundles that were located outside of hippocampus) and the number of functional hippocampal wires we recorded from. In the right column, the referencing and re-referencing scheme is indicated.

| Patient ID | # microwire bundles | # functional hippocampal wires | References |
| --- | --- | --- | --- |
| 01 | 4 (+ 2 removed due to artifacts) | 29 | Low impedance wires |
| 02 | 6 | 48 | High impedance wires |
| 03 | 6 | 42 | Ground, re-referenced to bundle mean |
| 04 | 4 | 32 | Low impedance wires |
| 05 | 5 (1) | 40 | Low impedance wires |
| 06 | 6 (2) | 48 | Low impedance wires |
| 07 | 5 | 39 | Low impedance wires |
| 08 | 2 | 16 | Low impedance wires |
| 09 | 2 | 16 | Low impedance wires |
| 10 | 2 | 16 | Low impedance wires |
| **Total:** | **42 (3)** | **326** |  |

**Table S7.** Task and recording information per patient, as well as the number of trials that were included in the analyses.

| Patient ID | Task version | # sessions | # correct encoding tr | # incorrect encoding tr | # correct retrieval tr | # incorrect retrieval tr |
| --- | --- | --- | --- | --- | --- | --- |
| 01 | 12. Image cue | 1 | 48 | 14 | 50 | 14 |
| 02 | 12. Image cue | 1 | 55 | 9 | 55 | 9 |
| 03 | 12. Image cue | 1 | 61 | 3 | 61 | 2 |
| 04 | 13. Verb cue | 2 | 99 | 27 | 101 | 20 |
| 05 | 13. Verb cue | 2 | 33 | 9 | 66 | 14 |
| 06 | 13. Verb cue | 3 | 90 | 31 | 92 | 31 |
| 07 | 13. Verb cue | 3 | 107 | 18 | 108 | 15 |
| 08 | 13. Verb cue | 2 | 116 | 12 | 116 | 12 |
| 09 | 13. Verb cue | 1 | 42 | 21 | 43 | 20 |
| 10 | 13. Verb cue | 2 | 103 | 25 | 103 | 25 |

### 2. Supplementary Figures S1 – S4

**Figure S1.**
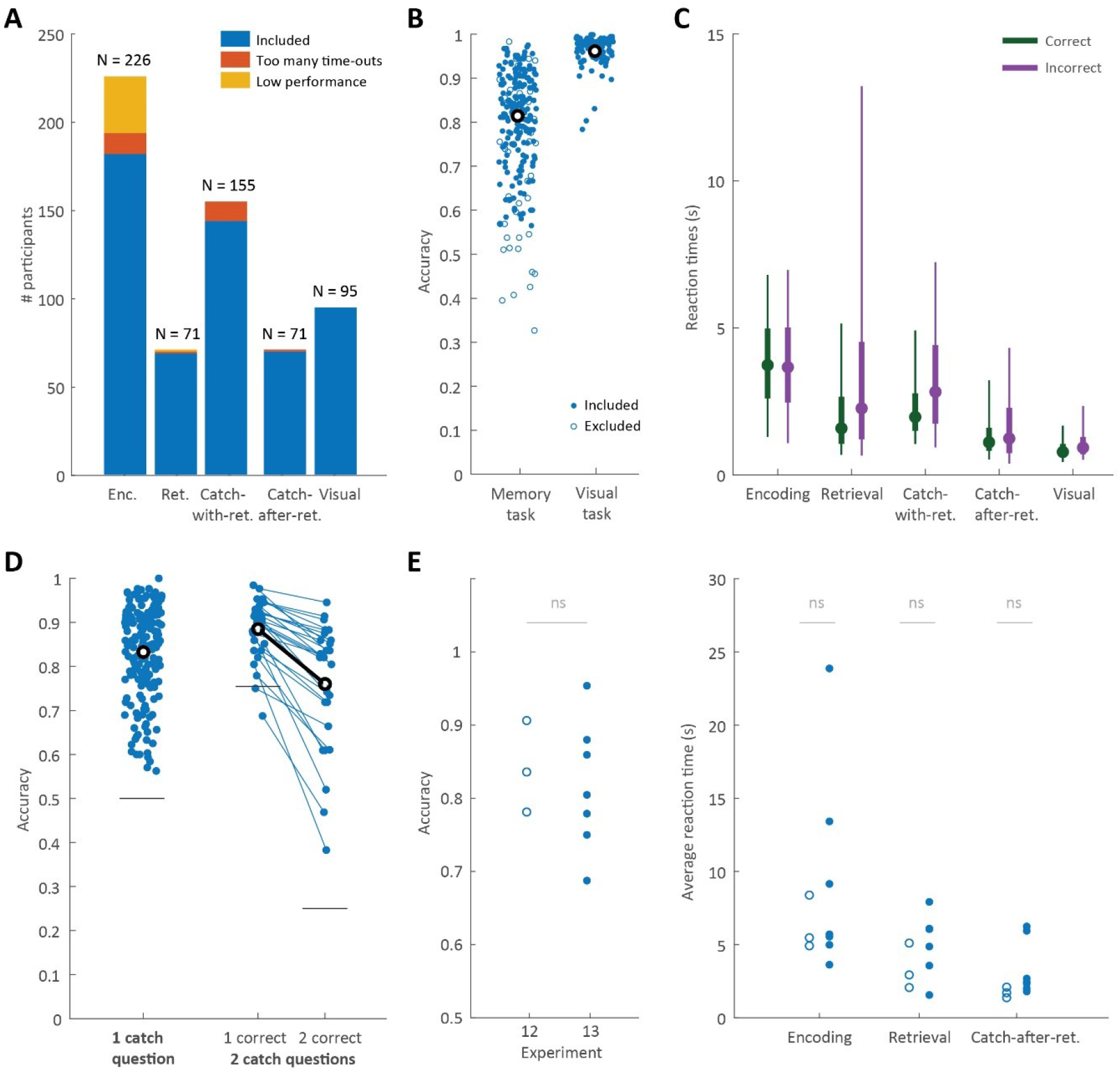
Descriptors of behavioral data. **A**: Number of included participants (blue) and the number of participants excluded for bad performance (yellow) or insufficient number of data points (orange) per task phase. Numbers above each bar indicate the total number of participants that were recorded. Enc. is Encoding; Ret. is Retrieval; Catch-with. is Catch-with-retrieval; **B**: Accuracy per participant on the memory and visual tasks. Included participants are indicated by filled circles, excluded participants by open circles; **C**: Box plots showing the reaction time distributions, across all included participants, for correct (green) and incorrect trials (purple), per task phase. Box plots indicate the 5, 25, 50 (circles), 75 and 95% boundaries; **D**: Fraction of correct trials for memory tasks with 1 catch question asked after retrieval (left), or two questions asked (middle and right). For participants answering two questions (connected dots), we show the fraction of trials with at least 1 question answered correctly (middle) and the fraction with two questions correct (right). Horizontal lines represent guessing level; **E**: Task performance (left) and reaction times (right) of the iEEG patients who participated in experiments 12 (open circles) and 13 (filled circles). Statistical comparison using a two-tailed t-test (α = 0.05); ns is not significant.

**Figure S2.**
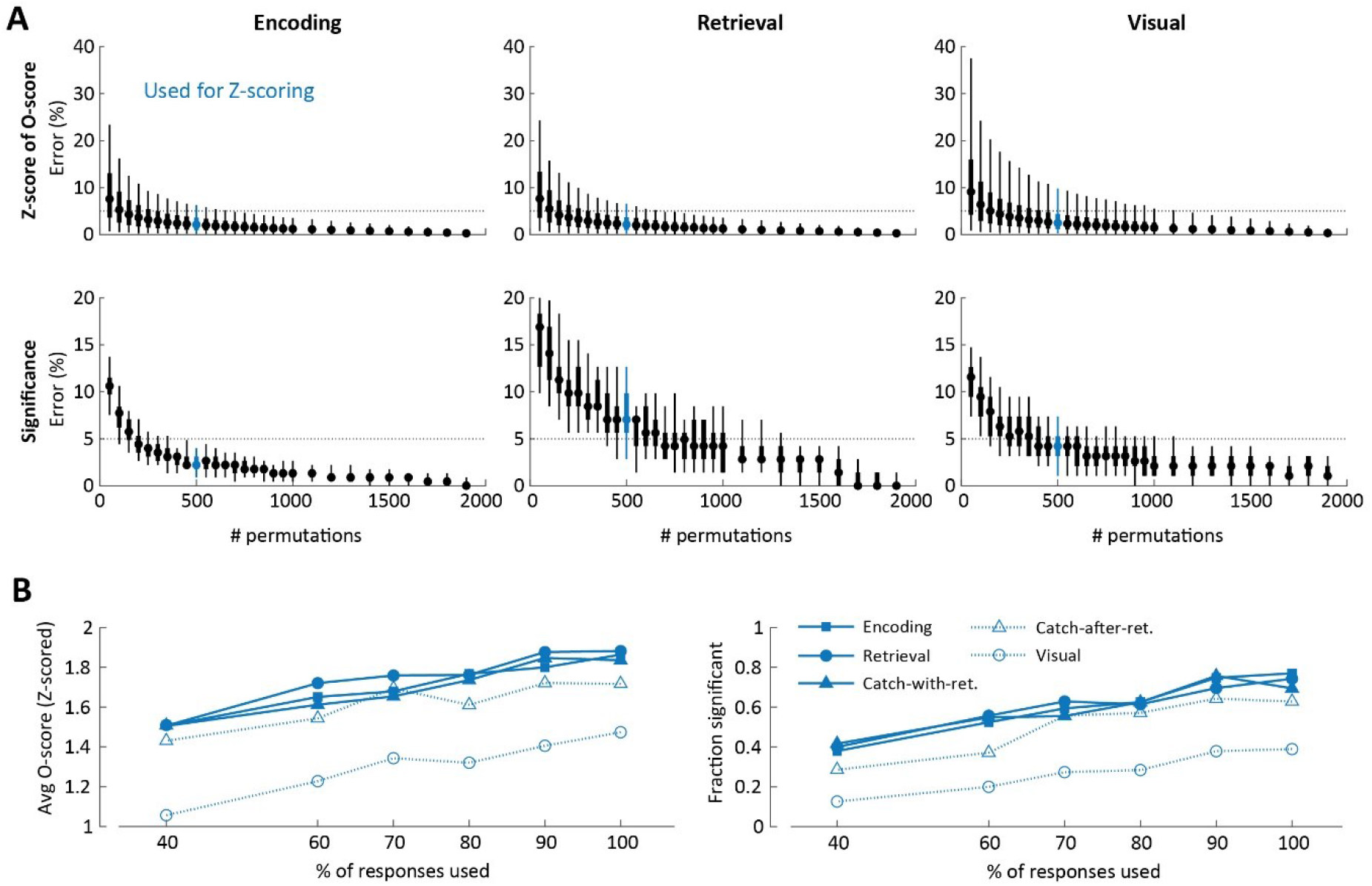
Validation of parameters of the behavioral analyses. **A**: To test the Z-scoring method for the O-score, we computed reference distributions with 2000 permutations for each participant for Encoding (left), Retrieval (middle) and Visual (right) task phases, and down-sampled the reference distribution to between 50 and 1900 permutations (x-axes), each repeated 50 times. We report the deviation (%) relative to the Z-scores obtained for 2000 permutations (top row) and the percentage difference in the number of participants marked as significant (bottom row). The number of permutations used throughout the manuscript (500 permutations), is indicated in blue. Horizontal dashed lines indicate a 5% error; **B**: Average Z-scored O-score (left) and fraction of participants with a significant O-score (right) as a function of the percentage of responses used to analyze the O-score. Reducing the percentage of responses included in the O-score analysis reduced the O-score and the fraction of significant participants, but this reduction affected all 5 task phases in a similar way. Task phases are indicated by line/symbol combinations: solid line with filled squares: Encoding; solid line with filled circles: Retrieval; solid line with filled triangle: Catch-with-retrieval; dashed line with open triangle: Catch-after-retrieval and dashed line with open circle: Visual.

**Figure S3.**
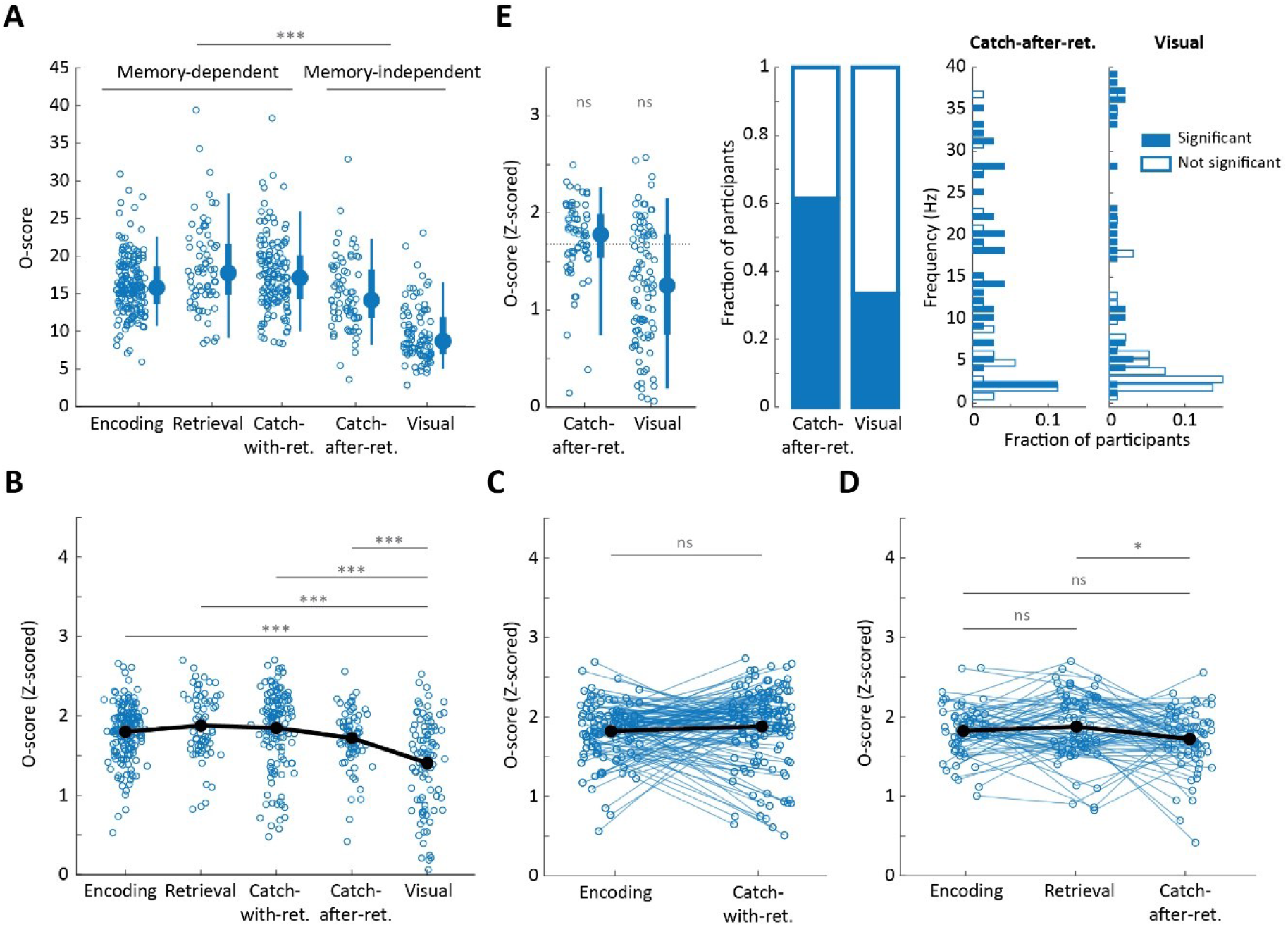
Additional O-score analyses. A: Raw O-scores per task phase. Each dot is one participant. Box plots (right) summarize the data across participants and show the 5, 25, 50 (circles), 75 and 95% boundaries; **B**: t-tests between Visual task phase and all other task phases (two-tailed, α = 0.05, Bonferroni corrected for 4 comparisons). Black line shows the mean O-scores; **C**: Paired t-test between task phases for memory task group 1 (two-tailed, α = 0.05). Connected dots are from one participant; **D**: Paired t-test between task phases for memory task group 2 (two-tailed, α = 0.05, Bonferroni corrected for 3 comparisons). Connected dots are from one participant; **E**: Z-scored O-scores, fraction of participants with significant O-scores and frequency distributions for Catch-after-retrieval and Visual task phases using a lenient lower frequency boundary. The lower frequency boundary was reduced from a period of 1/3 of the time series to 2 times the time series’ length (i.e. 6 times lower). Statistical comparison of the O-score as in the main text. ns: not significant; *: 0.05 ≥ p > 0.01; **: 0.001 ≥ p > 0.001; ***: p ≤ 0.0001.

**Figure S4 - part 1.**
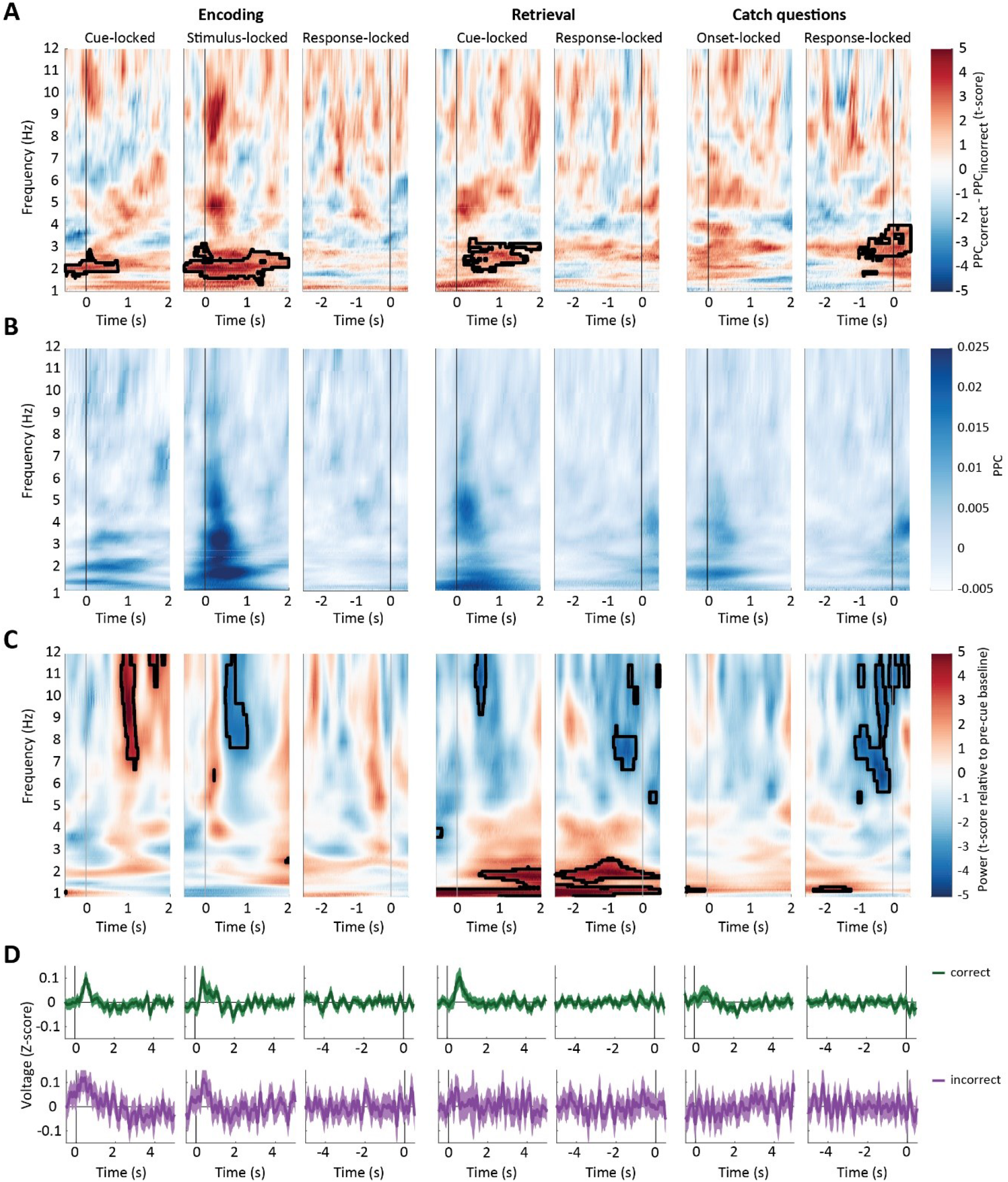
Supplementary data for Pairwise Phase Consistency (PPC) analyses from Figure 5. **A**: PPC for correct versus incorrect trials (t-scored, color-coded). Black outlines indicate significance (α = 0.05) compared to a reference distribution with permuted trial-labels; **B**: Averaged raw PPC values for correct trials across all subjects. Significant PPC increases (see Figure 5B) are accompanied by increases in raw PPC; **C**: Average power changes relative to baseline for correct trials across all subjects; **D**: Event-related potentials (ERPs) for correct (green, top row) and incorrect trials (purple, bottom row). Voltage traces were Z-scored per trial relative to pre-cue baseline and averaged across trials, channels and subjects. Solid lines show the smoothed (50 ms kernel) mean voltage and shaded areas indicate the standard error of the mean. All panels show data from encoding trials (left, cue-, stimulus- and response-locked), retrieval trials (middle, cue- and response-locked) and catch questions (onset- and response-locked), with vertical black lines indicating t = 0 s.

**Figure S4 - part 2.**
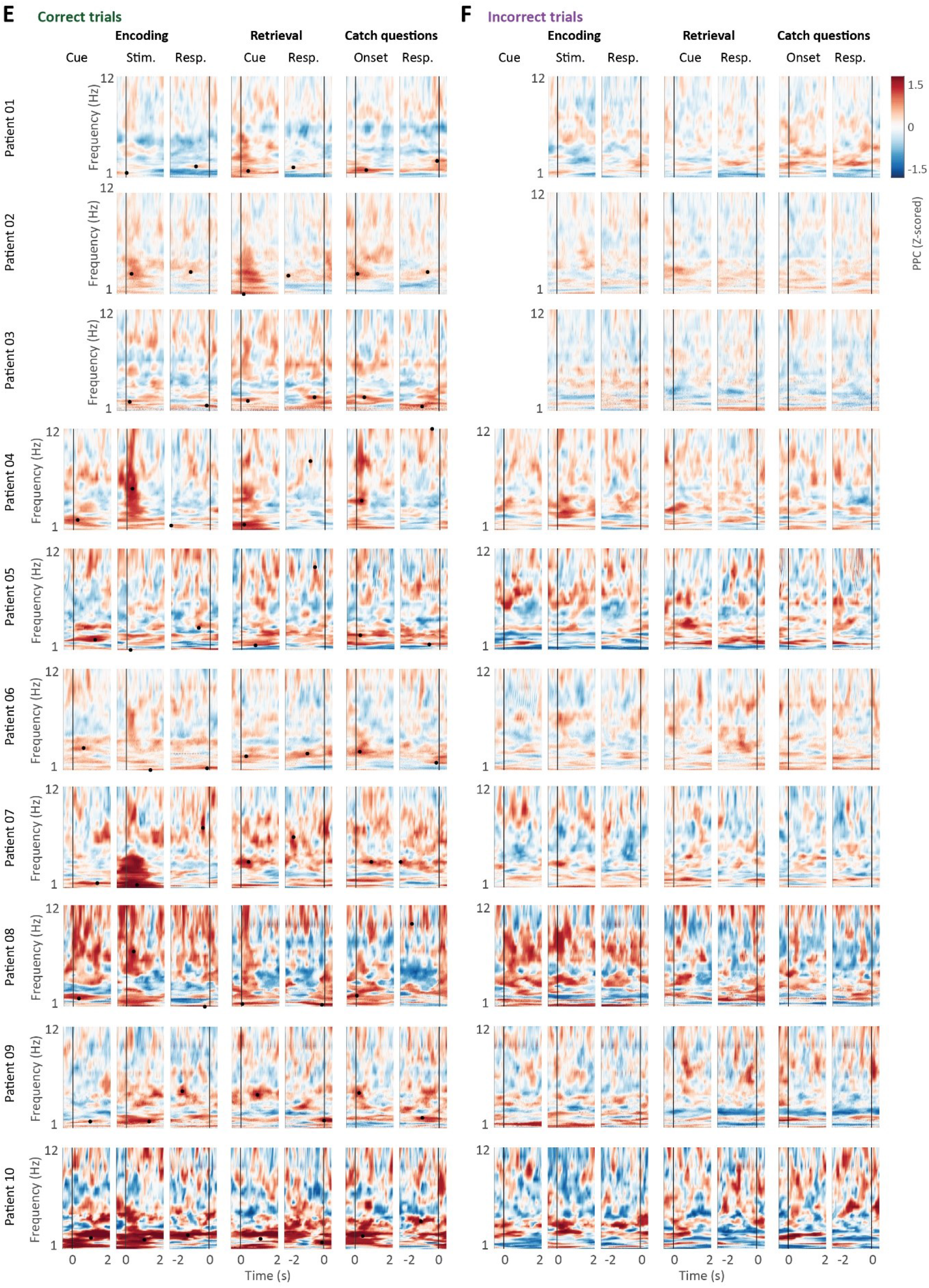
Average Pairwise Phase Consistency (PPC), Z-scored relative to pre-cue baseline (color-coded) for each of the 10 intracranial EEG patients for correct (**E**) and incorrect trials (**F**), and locked to cue onset, stimulus onset and response for encoding trials, cue onset and response for retrieval trials, as well as for catch question onset and response.

**Figure S4 - part 3.**
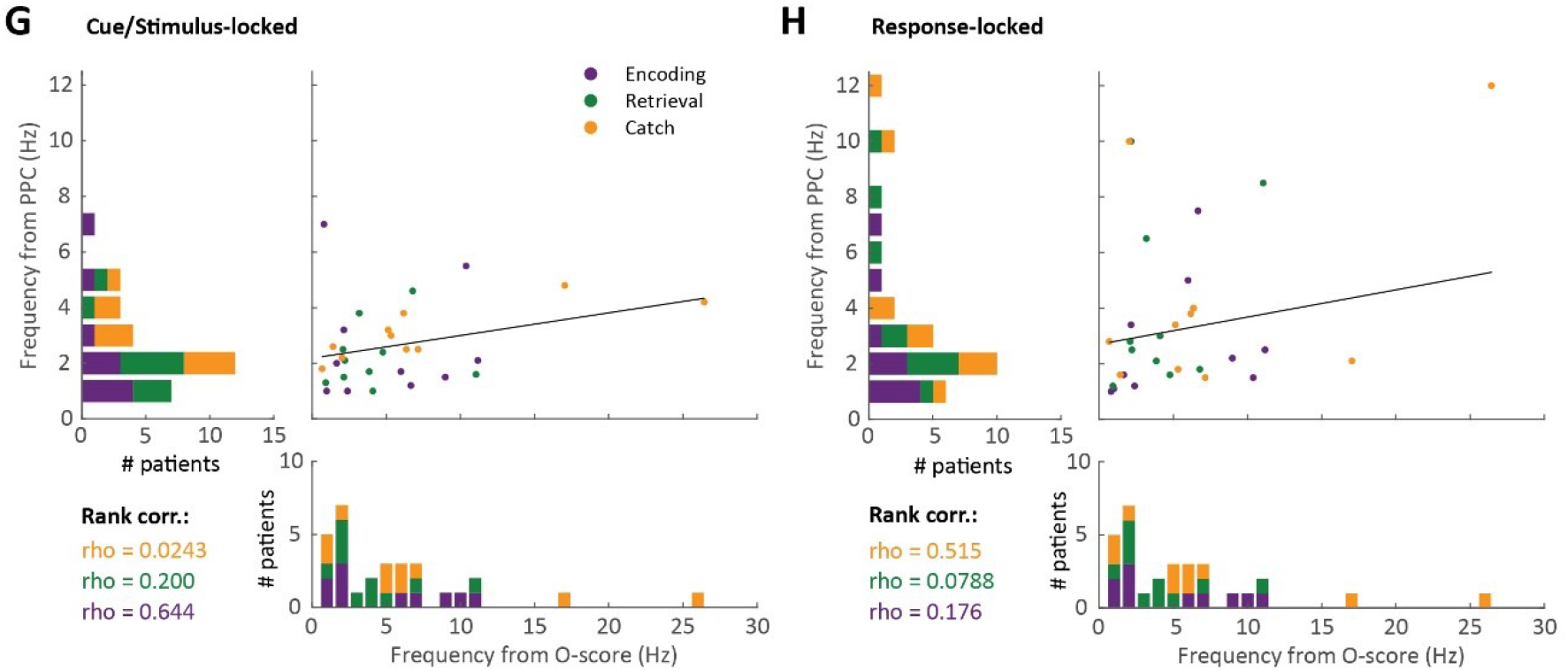
Supplementary data to phase analyses in Figure 5 and comparison of O-score analyses with peak PPC frequencies. **G&H**: Scatter plots and histograms of the peak frequencies identified by the O-score (horizontal axes) and the peak PPC (vertical axes) for cue/stimulus locked analyses (G) and response-locked analyses (H) of the PPC, and for encoding (purple), retrieval (green) and catch (yellow) phases separately. The Spearman rank correlation is given for each task phase separately in the same color code.

### 3. Supplementary Methods: Simulated data for validation of O-score method

To test the applicability and performance of the O-score method for our data we ran a series of simulations. In these simulations, we produced model data with and without oscillations, i.e. with a known ground truth, and computed O-scores in the same way as reported for the behavioral data. This allowed us to test whether the O-score method is able to detect oscillations of different strengths in a noisy dataset, as well as reject data with no oscillation included. In addition, it allowed us to compare the O-score’s ability to detect oscillations for datasets with different characteristics, such as average response times, with each other. To this end, we modelled data that matched the response distributions, the number of responses and the number of included participants from the encoding and visual task phases.

Response data were simulated as a Poisson process, i.e.:

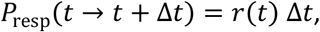

with *P*_resp_ the chance of a response occurring at the time interval *t* to *t* + Δ*t*, *r*(*t*) the ‘firing rate’ at time *t* and Δ*t* the time step, set to Δ*t* = 0.0005 s. The firing rate function *r*(*t*) was modelled as:

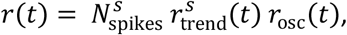

i.e. as the product of the total number of spikes *N*_spikes_, an overall trend function *r*_trend_(*t*) and an oscillatory function *r*_osc_(*t*), of which the former two factors were varied between model participants *S*, introducing noise in the data set. The total number of spikes was drawn from a normal distribution, the mean and SD of which were matched by the behavioral data (Table S3) and with a minimum of 10 spikes.

For the visual task phase, the firing rate function *r*_trend_(*t*) was modelled using a gamma probability density function, while the trend for the encoding task phase was modelled using a normal probability density function and the retrieval task phase with a lognormal distribution. The length of the time series was varied between model participants and was drawn from a uniform distribution. The parameters for the probability density functions are given in Table S8.

To introduce the oscillation, we used:

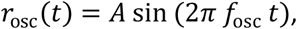

with *f*_osc_ the frequency of the oscillation, which was set to 2.5, 5, 7.5, 10 and 15 Hz, and *A* the amplitude, varying between 0 (i.e. no oscillation) and 1 (i.e. 100% modulation of response rate within each period), in steps of 0.1. The results for a range of *f*_osc_ and *A* is given for both model task phases in Figure S5.

**Table S8.** Parameters of the model data. See Supplementary Methods for details.

| Task phase: | # model participants | # spikes Mean ± SD: | $r_{\text{trend}}(t)$: | | | | | |
| --- | --- | --- | --- | --- | --- | --- | --- | --- |
|  |  |  | Pdf function | Parameter 1 |  | Parameter 2 |  | Total time (s) |
| Encoding | 190 | 66 ± 34 | Normal | Mean: | 1.5-2.5 | St dev: | 2.5-3.5 | 4-12 s |
| Retrieval | 70 | 151 ± 54 | Lognormal | Mean: | 0-1 | St dev: | 1-1.5 | 4-12 s |
| Visual | 95 | 215 ± 54 | Gamma | Shape: | 1-2 | Scale: | 0.25-0.5 | 1.5-4.5 s |

### 4. Supplementary Results: Validation of O-score method

We simulated data resembling the encoding, retrieval and visual task phases in terms of the number of participants, reaction time distributions and number of responses per participant, with various levels of oscillatory modulation. Some example simulations are given in Figure S5A. Similar to the analyses on the observed data, we computed the O-score for each simulated participant. To aid comparison with Figure 3 of the main text, we report the distributions of O-scores across participants (mean and standard deviation given in the top rows of Figure S5 B-D), and the number of participants reaching the significance threshold (bottom rows of Figure S5 B-D). We also computed second level statistics and report whether the results are significant at α = 0.01. Finally, we computed frequency histograms across significant O-scores (middle rows of Figure S4 B-D).

The results in Figure S5 provide us with two important validations. Firstly, we observe that O-scores were low and did not reach significance when no or weak oscillatory modulations were applied, and this was the case for each of the 3 task phases. This suggests that the O-score analysis is not likely to provide spurious results when an oscillation is not present or is too weak to be detected.

Conversely, for stronger oscillations, O-scores became significant for each of the three task phases, and the fraction of participants with significant O-scores went up markedly. Importantly, when the O-score analysis indicated significance at the population level, then the peak frequencies were overwhelmingly detected at the correct frequency (red dotted lines in Figure S5 indicate a 2 Hz frequency band around the ground truth frequency).

The oscillation amplitude at which the significance threshold was reached differed per task phase, with a low threshold of 20-30% modulation sufficient for Retrieval and a high 50-60% required for Encoding. This differentiation is expected given the large differences in response density between the task phases, with extremely low response densities for Encoding (i.e. few responses over a long timeframe). We note that O-scores remained non-significant and the fraction of significant participants was low when the oscillation frequency was expected to be undetectable given the characteristics of the data set, again suggesting spurious results are unlikely. This occurred for the 15 Hz oscillation for Encoding, due to low response density and corresponding reduction in the upper frequency bound. O-scores also dropped for low oscillation frequencies for the Visual task, due to the short response timeframe and corresponding increase in lower frequency bound. The latter drop in O-score could be ‘rescued’ by increasing the maximum detectible oscillation period (i.e. minimum frequency) for the standard 1/3^rd^ of the timeframe to 2 times the response timeframe (yellow Figure S5D), confirming the validity of the analyses shown in Figure 3E.

**Figure S5.**
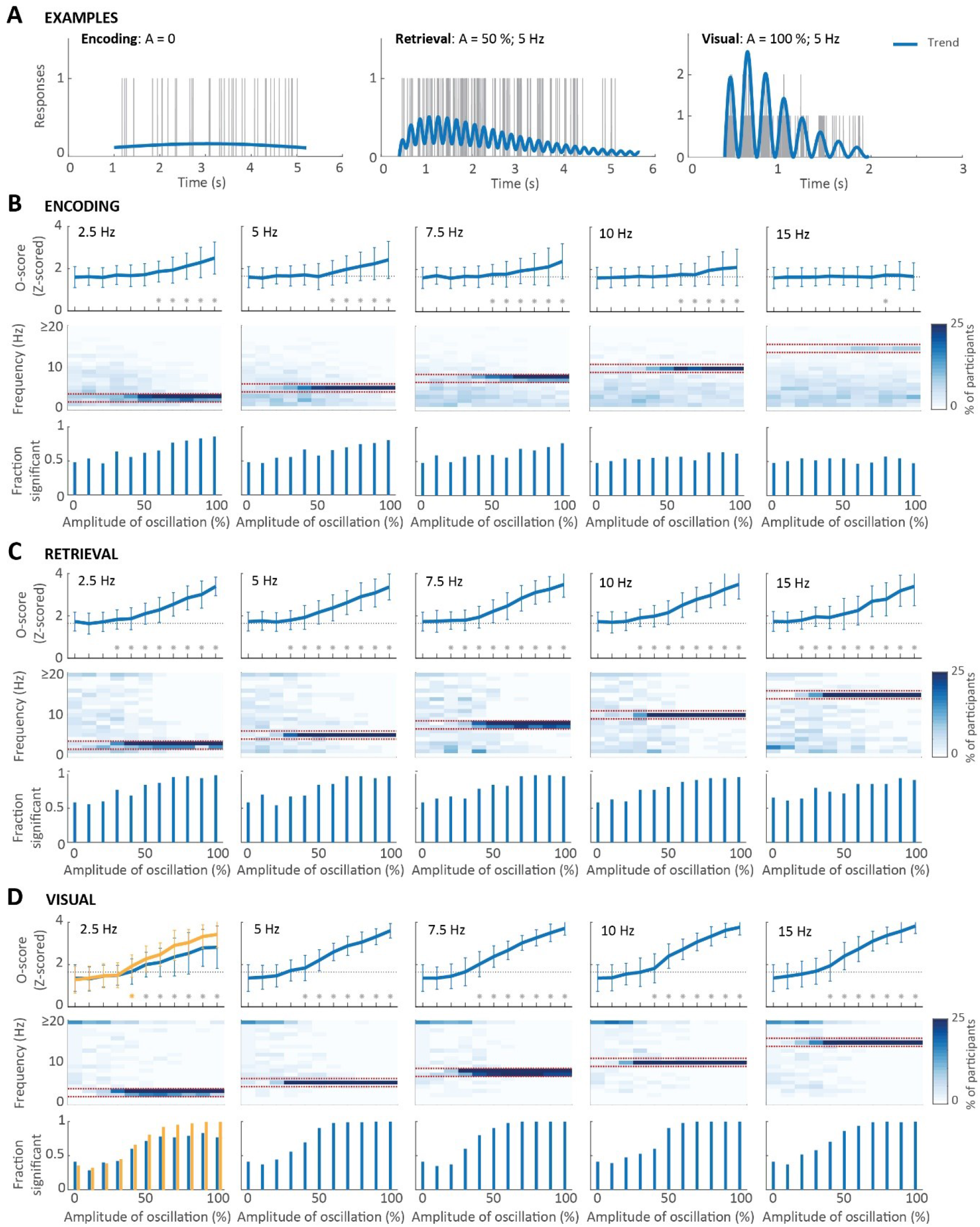
O-scores for simulated data for Encoding (B), Retrieval (C) and Visual (D), covering a range of oscillation amplitudes (x-axes; 0 % = no oscillation; 100 % = complete modulation), for 5 frequencies (left to right): 2.5, 5, 7.5, 10 and 15 Hz. **A**: Example traces, with trend curves in blue and response traces in grey. **B-D**: For each task phase, we show the Z-scored O-score (top row, mean ± standard deviation, * indicate significance for population at α = 0.01), frequency histogram per amplitude (middle row, color-coded, note: y-axis truncated at 20 Hz) and fraction of participants with significant O-scores (bottom row). In D, results of an additional simulation are shown for 2.5 Hz, were the lower frequency bound for O-score computation was reduced by a factor 6 (yellow).

